# Proteome allocation and the evolution of metabolic cross-feeding

**DOI:** 10.1101/2021.12.17.473181

**Authors:** Florian J. F. Labourel, Vincent Daubin, Frédéric Menu, Etienne Rajon

**Author notes:** **Data availability:** All the scripts generated for this project are available at: https://github.com/FloTuzoLab/Scripts-Evolution-CF-proteome-allocation. **Competing interests:** The authors declare no competing interests.

## Abstract

Metabolic cross-feeding (MCF) is a widespread type of ecological interaction where organisms share nutrients. In a common instance of MCF, an organism incompletely metabolises sugars and releases metabolites that are used by another as a carbon source to produce energy. Why would the former waste edible food, and why does this preferentially occur at specific locations in the sugar metabolic pathway (acetate and glycerol are preferentially exchanged) have challenged evolutionary theory for decades. Addressing these questions requires to model the cellular features involved; to this end, we built an explicit model of metabolic reactions, including their enzyme-driven catalysis and the cellular constraints acting on the proteome that may incur a cost to expressing all enzymes along a pathway. After showing that cells should in principle prioritise upstream reactions when metabolites are restrained inside the cell, we investigate how the diffusivity of these metabolites may trigger the emergence of MCF in a population. We find that the occurrence of MCF is rare and requires that an intermediate metabolite be extremely diffusive: indeed, up to high membrane permeability coefficients, the expected evolutionary outcome is not a diversification that resembles MCF but a single genotype that instead overexpresses downstream enzymes. Only at very high levels of membrane permeability and under distinctive sets of parameters should the population diversify and MCF evolve. These results help understand the origins of simple microbial communities, and may later be extended to investigate how evolution has progressively built up today’s extremely diverse communities.

**Significance statement:** Can two species thrive on a single energetic resource? While the competitive exclusion principle predicts that one in the pair should go extinct, it may occur that an organism releases partly metabolised molecules in the environment, securing an ecological niche for a second organism in a specialisation process called metabolic cross-feeding. Here we investigate how evolution may favor the waste of a useful resource using a model that considers how a cell packed with proteins may be less efficient, hence favoring a shortening of metabolic pathways in order to reduce cell packing. Our model indicates that such specialisation only occurs under restricted conditions. Incidentally, this makes the signatures of cross-feeding, such as which metabolites are preferentially involved, quite predictable.

## Introduction

Genetic diversification [1, 2] may occur when different ecological niches are encountered [3–5], for instance when different carbon sources are available in the environment [6, 7]. What may at first glance sound puzzling – why not using all the available nutrients? – finds an explanation in physiological and environmental constraints or even absolute incompatibilities that make specialists of each resource outperform generalists [8–13]. Even more bewildering is the observation that diversification occurs in the homogeneous presence of a single energetic resource [14–17]. One finds a clear example in chemostats or controlled experimental systems in which glucose is continuously injected, where glucose consumers may evolve that release metabolites for others to use as carbon sources [14, 18, 19]. This unidirectional by-product process is a form of metabolite cross-feeding [20, 21], and its evolutionary underpinnings are still blurry [20, 22].

In particular, the reasons why specific metabolites are more likely involved in cross-feeding remain unclear. Indeed, a large number of metabolites produced by a glucose-reliant organism may constitute viable single carbon sources for others [for example, a glucose-reliant strain of *Escherichia coli* can theoretically produce up to 58 useful metabolites, 22]. Yet only two metabolites are commonly reported – from experiments – as being traded in such cross-feeding interactions, namely acetate and glycerol [14, 19, 23, 24]. In line with the fact that some of these lineages may have been predisposed to use these metabolites [23], San-Roman and Wagner [25] have hypothesized that this preferential evolution could be due to shorter mutational paths. Accordingly, modifying the metabolic network to produce these interacting strains would require fewer and less destabilising mutations and could thence arise more readily. But their conclusion is that acetate or glycerol trades are no more likely than others to appear through mutation, making the mystery about their involvement – and potential predispositions of strains – in the evolution of metabolic cross-feeding even deeper.

Adaptation is often incomprehensible without considering the ability of an organism to perform a task as being dependent on internal constraints [26–29]. For example, fully expressing all enzymes along a metabolic pathway may incur a fitness cost [30, 31], to such an extent that sacrificing a part of a pathway becomes beneficial [32, 33]. Cells whose cytoplasm gets crowded with proteins actually pay a two-fold cost [34]. First, producing enzymes incurs a direct energetic expense, approximately proportional to the sum of enzyme concentrations in the cell [35–37]. Second, cell packing eventually compromises the diffusion of proteins, thereby hindering metabolic efficiency [38, 39].

In a previous study, we have shown that the evolution of enzyme kinetic parameters and concentrations is contingent on their competition with other processes for their substrate [34]. One of these competing processes may be leakage through the cell membrane, such that highly diffusive metabolites should be processed by more efficient or concentrated enzymes. The combination of this requirement for high concentrations, and the cost of an abundant proteome, could make these metabolites the preferential breakpoints in a metabolic pathway. Very few metabolites can diffuse through membranes, either because of their size or due to their electronic properties [40]. Such diffusion may be direct, as is the case for glycerol, or indirect when a non-diffusive metabolite can spontaneously transform into a diffusive one, as is the case with acetate [41, 42].

In this work – see Figure 1 for an overview of the metabolic model – we first determine how cells should allocate their proteome when metabolite diffusion is limited. We find that upstream reactions should be favored when selective pressures are similar along the pathway. We then assess the hypothesis that cross-feeding evolves in response to the high diffusion rate of an intermediate metabolite, due to the incurred selection on downstream enzymes. Indeed, a genotype producing the diffusive metabolite must also be very efficient at metabolising it to prevent its loss, and thus pays a high cost over-expressing downstream enzymes that, at some point, may become insurmountable. Interestingly, a second genotype feeding on the intermediate metabolite and only expressing downstream enzymes would thrive in this context where the metabolite has become a resource. This is because, on top of saving on the expression of upstream enzymes, extensive over-expression downstream is no longer a requirement as the high permeability coefficient of the metabolite actually helps its uptake. Yet, the overall evolutionary process must be continuous, instead of the schematic two-steps sequence presented here, making it difficult to predict when the evolution of cross-feeding should occur.

**Figure 1.**
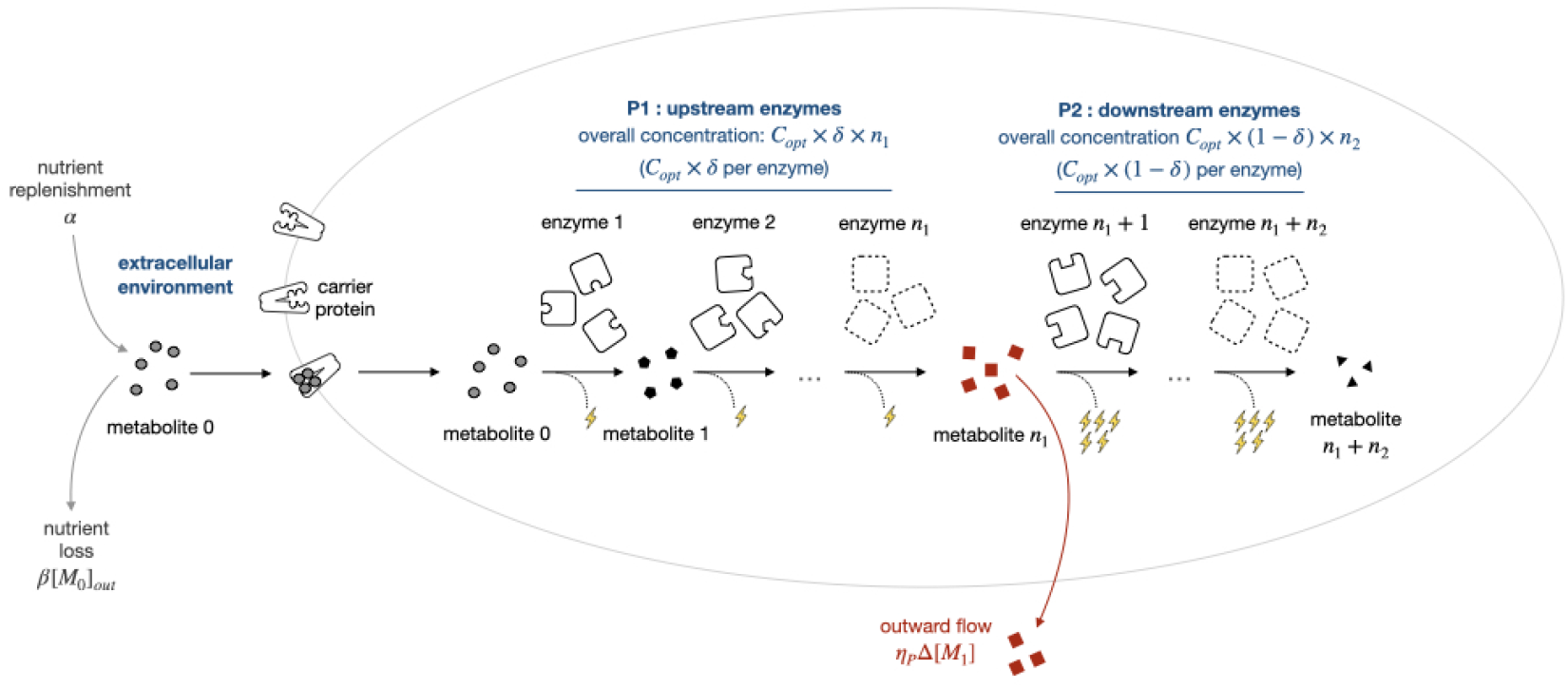
Overview of the model: the pathway is initiated by a carrier protein and comprised of *n*_1_ upstream enzymes defining the sub-pathway *P*_1_ and *n*_2_ downstream enzymes defining the sub-pathway *P*_2_. Allocation of the proteome is driven by two parameters: (i) *C*_*opt*_ coincides with the adaptive proteome fraction dedicated to one of the reaction of the full pathway *P*_1_ + *P*_2_ when all enzyme concentrations are identical (the case of no differential allocation – see section Optimal metabolic allocation); (ii) the parameter *δ* (*resp*. 1 − *δ*) corresponds to the fraction dedicated to the first sub-pathway *P*_1_ (*resp*.*P*_2_). The extracellular dynamic of the nutrient (metabolite 0) is based on a constant replenishment-degradation process – see Materials and Methods section. Within the pathway, each reaction follows Briggs Haldane kinetics (with or without reversibility) where enzyme efficiency is constant and studied as a parameter. These reactions each provide a fitness yield, shown above as lightning, proportional to the amount of product produced and to the specific yield of energy gain set for reactions (1 energy unit per reaction per nutrient for upstream enzymes and 5 for downstream enzymes in the above example) – see section on overexpression. Fitness is simultaneously impeded by the cost of expression – production and crowding – and, depending on the subsection again, by the toxicity resulting from the total concentration of metabolites. The metabolite produced at the end of the upstream pathway *P*_1_ is susceptible to leak through the membrane according to a permeability *η*_*P*_ – studied as a parameter – and the gradient of concentration between the extra and intracellular environments. Once in the extracellular environment, it is available to other cells but may also be degraded (like metabolite 0)

In order to embrace this continuity, we use Adaptive Dynamics [43, 44] to model the evolution of the pattern of enzyme expression along the pathway. This framework is particularly suited to model the complex ecological interactions that may arise as the genotype(s) composing the population shape their environment [32, 45], by controlling the equilibrium frequencies of both the nutrient and the diffusing metabolite. We find that MCF sometimes evolves, characterised by an evolutionary diversification giving rise to the coexistence between a genotype only expressing the enzymes upstream the diffusing metabolite, and another one expressing the enzymes down-stream. This occurs at very high membrane permeability levels of the diffusing metabolite, only compatible with diffusion rates reported for acetate or glycerol. We also find that MCF requires moderate to high levels regarding the intracellular selective constraints acting on metabolites along the pathway.

## Results

### Optimal metabolic allocation and cell constraints

#### Evolution of the overall expression of metabolic enzymes

We first assume that all enzymes have an equal concentration and consider its evolution. Increasing concentration enhances the efficiency of catalysis and thus the production of energy, but with diminishing returns [34, 46]. It also incurs costs, firstly due to the actual energy cost of making proteins, and secondly because high protein concentrations in the cell decrease the efficiency of reactions due to cell packing. The former is captured in our model by a linear cost inflicted to extra production, and the latter through a penalty on *k*_*f*_, whose effect has been estimated [39, 47] and modelled in previous studies [38] – [see Model and 34].

To approach the case of reactions involved in the carbon cycle [48], we consider a pathway comprised of 40 reactions and initiated by facilitated diffusion through a transporter [49], where energy is produced at each step in the process – see Figure 1 – and assessed the effect of reaction yield – see SM Figure S3. Reactions follow irreversible Briggs-Haldane kinetics [50], with kinetic constants set by default half an order of magnitude higher than the median observed for enzymes involved in the central metabolism [51] (*k*_*f*_ = 10^6.25^*M*^−1^*s*^−1^, *k*_*cat*_ = 10^2.25^*s*^−1^) – see SM Table S2 for the full set of parameters we used; besides, we also consider the case where reactions are reversible see SM Text S1.2. Setting these efficiencies above the average found for enzymes involved in central carbon metabolism is conservative as it diminishes the selective pressure that promotes higher enzyme expression, and thus ensures that we are not examining an implausible range of values (*eg*. because the whole set would be biased by some enzymes that need not be efficient due to specific metabolic features). The local context of reactions – including reaction reversibility and metabolite toxicity – may also affect metabolic efficiency, as captured by a linear degradation rate of each metabolite *η* in this instance of the model [34] – see SM Text S1.2 for more complex selective pressures. Nutrients are added to the environment at a constant rate *α* and degraded at a linear rate *β* – the latter also applying to metabolites released by cells in the medium (see below) whose levels are such that without cells, substrate concentration equilibriates at 1*M*.

For all combinations of the parameters above considered, the evolutionarily expected concentrations of the 40 enzymes in the pathway sum up to 15 − 20 % of the whole proteome (see SM section Text S1). As we have shown previously, adaptive outcomes depend on a complex interaction between cell constraints, enzymes concentrations and kinetic efficiencies that cannot be captured through the influence of classical experimenters’ parameters (such as *K*_*M*_ or the activity constant *k*_*cat*_[*E*_*tot*_]*/K*_*M*_, for instance) alone [34]. The highest fraction is self-evidently obtained in conditions where selection for the rates of reactions is acute, such that increasing concentration becomes beneficial, to a certain extent, despite amplifying intracellular crowding and production costs. These predictions are consistent with estimates among unicellular species [48]: in most cases, enzymes involved in the carbon cycle constitute approximately 20 % of the proteome. In the remaining of this study, the overall concentration of enzymes in the pathway is considered fixed at its evolutionary expectation by default, *i*.*e*. that obtained for the specific combination of parameters studied when assuming an identical allocation all along the pathway.

#### Overexpression in upstream reactions

We then studied how cells should distribute enzyme expression along a pathway split into two parts of equal lengths. This is a proteome allocation problem, since the overall concentration is fixed to an (adaptive) optimum as just described; we study the evolution of the part of this overall concentration allocated to the first half *P*_1_ of the metabolic pathway, *δ*, which we assume can change by mutation in the range [0, 1] – *δ* = 0 coinciding with no investment in *P*_1_ such that all of the resources are allocated to *P*_2_, while *δ* = 1 corresponds to all resources being allocated to *P*_1_ (and none to *P*_2_).

The evolution of *δ* is modelled using adaptive dynamics [32, 43, 44], as is appropriate when the fate of a mutant can depend on the environment shaped by one (or several) resident genotype(s). Here the resident strategy may impact the equilibrium concentrations of the nutrient and of the metabolites produced along the pathway. Nonetheless at this stage, each metabolite is considered to be unable to diffuse across the membrane, such that their concentration in the environment remains zero. Therefore in this case, the evolutionary outcome is always a single allocation strategy *δ*, as exemplified in Fig. 2. In this figure, whatever the initial resident strategy in place in the population, evolutionary trajectories will converge to *δ* ≈ 0.6, and, once in place in the population (as resident) this latter strategy will be stable against the invasion by mutants with any other *δ*. These features make an enhanced expression of upstream reactions – with *δ* ≈ 0.6 – the evolutionarily expected outcome, also described as a convergent stable strategy – CSS, hereafter.

**Figure 2.**
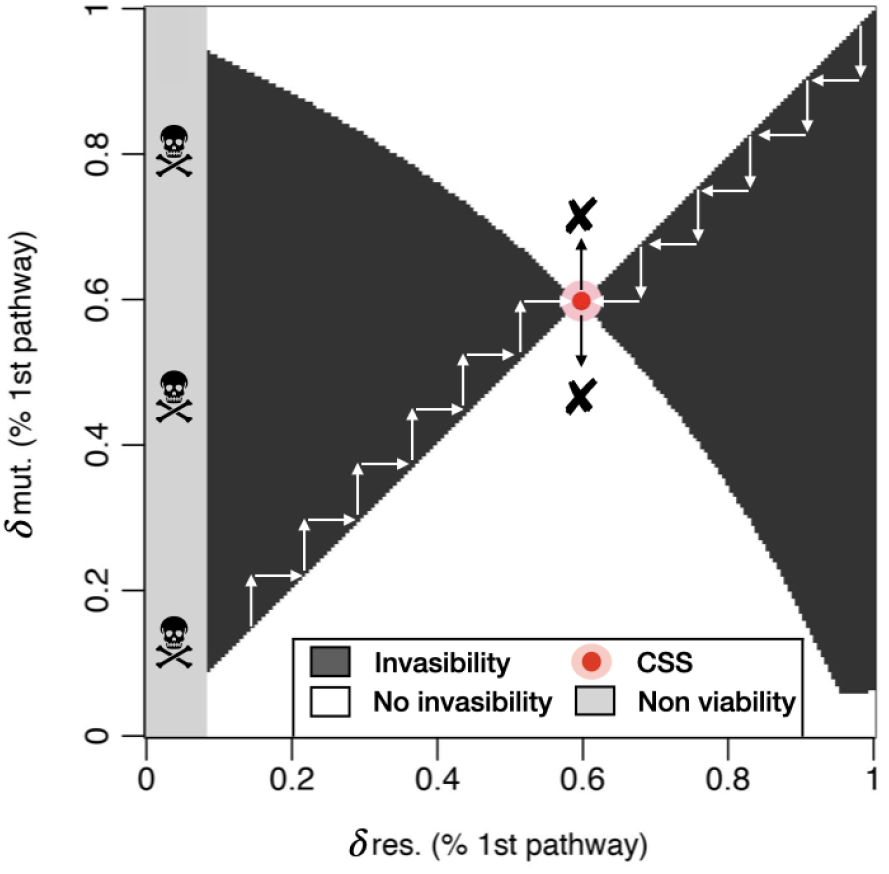
Example of the outcome of Adaptive Dynamics shown through a pairwise invasibility plot – PIP, here-after – where black (resp. white) areas stand for positive (resp.negative) invasive fitness. The grey area represents an area where no strategy is viable because it cannot produce enough energy to reach the level set as corresponding to the demographic steady-state even when only one individual is present. When a mutant arrives in the population of resident strategies, it takes the place of the resident strategy if its invasion fitness is positive, a process represented by the white arrows. Here, below the CSS, mutants with higher values (black area above the left lower corner to right upper corner bisector) than the resident invades and evolution pushes resident to converge towards the CSS, while above the CSS, it is the other way around. This PIP shows how cells should spread their content, according to the respective fractions *δ* and 1 − *δ*, between upstream and downstream enzymes and exemplifies that under the specific set of parameters chosen, they should prioritise upstream reactions since *δ* > 0.5, as was generally found – see text for more details.

In the presence of degradation, the adaptive investment in the first part of the pathway is generally above 0.5 : remarkably, *δ* evolves to more than 0.6 even at very low degradation rates where the resulting loss in metabolites is less than 1% along the pathway (see Table S3 of SM for an analysis of the influence of degradation on the loss of metabolites and Figure S5 of SM for more details). One factor that pushes upstream expression towards higher adaptive values is the selective pressure imposed by transporters – see Figure S5-B of SM for the influence of transporters. Yet, the process also holds when this influence is set apart. A plausible explanation is that within an irreversible pathway as the one modelled, upstream enzymes not only concur to fitness directly through the energy generated by their respective reaction, but also through their indirect contributions to downstream reactions – see SM Text S2 and Figure S6 for the analysis of this phenomenon through a more tractable model showing that the optimal strategy should always be to prioritise upstream reactions. This is also consistent with the fact that this unequal allocation holds when downstream reactions produce far more energy than their upstream counterparts so that upstream reactions contribute mostly indirectly to fitness (and is significantly heightened in the opposite case): even when increasing the yield of the reactions in the second half of the pathway tenfold as described for the carbon cycle [52], *δ* remains close to 0.6. Besides, the irreversible loss of metabolites caused by an increase in the degradation rate increases the asymmetry in fitness contributions further and thereby tends to increase the adaptive ratio of upstream to downstream enzyme expression – see Figures S5 and S15 of SM.

Such asymmetries in fitness contributions might help explain why enzymes catalysing reactions more upstream tend to face stronger selection [53–55]. Overexpression of upstream enzymes can nonetheless be – at least, partly – counteracted by selection for homogeneity in metabolite concentrations, as is the case when toxicity is high (Figure S7-B of SM) and equally spread, and it also depends on reactions reversibility, including that of the transporter, inasmuch as reversibility also shifts the balance between direct and indirect contributions of each subpathway: if downstream reactions are more reversible than upstream ones, cells should prioritise them, and vice versa – see Figure S7-A of SM. Be that as it may, considering realistic combinations of these pressures – moderate toxicity and degradation rates as well as the average reversibility found in central carbon metabolism [56] tends to corroborate the need for upstream overexpression, as shown in Section Text S2.2 of SM. One constraint that can reshu?e the deck of cards is the permeability of the cell membrane, which was hitherto not considered.

### Membrane permeability and cross-feeding

#### Membrane permeability impacts proteome allocation

Membranes are only permeable to a few metabolites, owing to their unique chemical features [40]. In our model, we allowed the metabolite produced by the last reaction of the first half of the metabolic pathway to diffuse – see the model overview in Figure 1 and the Model section for details – with permeability rates ranging from *P* = 10^−12^ dm.s^−1^ to 10^−4^ dm.s^−1^, in line with empirical estimates [40]. Metabolites making their way across the membrane have two consequences. First, they are lost for the cell that has produced them, which may act as a selective pressure for limiting diffusion. Second, they may accumulate in the external environment and be available to other individuals as a resource.

A cell has little leverage to limit outward diffusion; the most obvious solution is to use the metabolite before it is lost, which in our model is possible through an increase in the concentration of enzymes acting on the second part of the pathway. This is indeed what happens: the optimal allocation shifts from a higher concentration in the first part of the pathway – owing to the aforementioned factors – to a higher concentration in the second part as permeability increases, as shown for instance in Fig. 3 in the case of a low degradation rate (grey dots). The results presented in Fig. 3 are for proteins with kinetic parameters being slightly higher than the median reported for enzymes in the carbohydrate and energy pathway [51]. Higher efficiencies consistently produce a qualitatively similar result of a downward shift in allocation to the first part of the metabolic pathway, *P*_1_, as permeability increases (see Figure S12 in SM). This outcome is also robust to the possible existence of different energetic yields along a pathway – see Figure S8 for the influence of yield on the optimal allocation – and most often holds when introducing other mechanistic constraints – see Figures S13 and S14.

**Figure 3.**
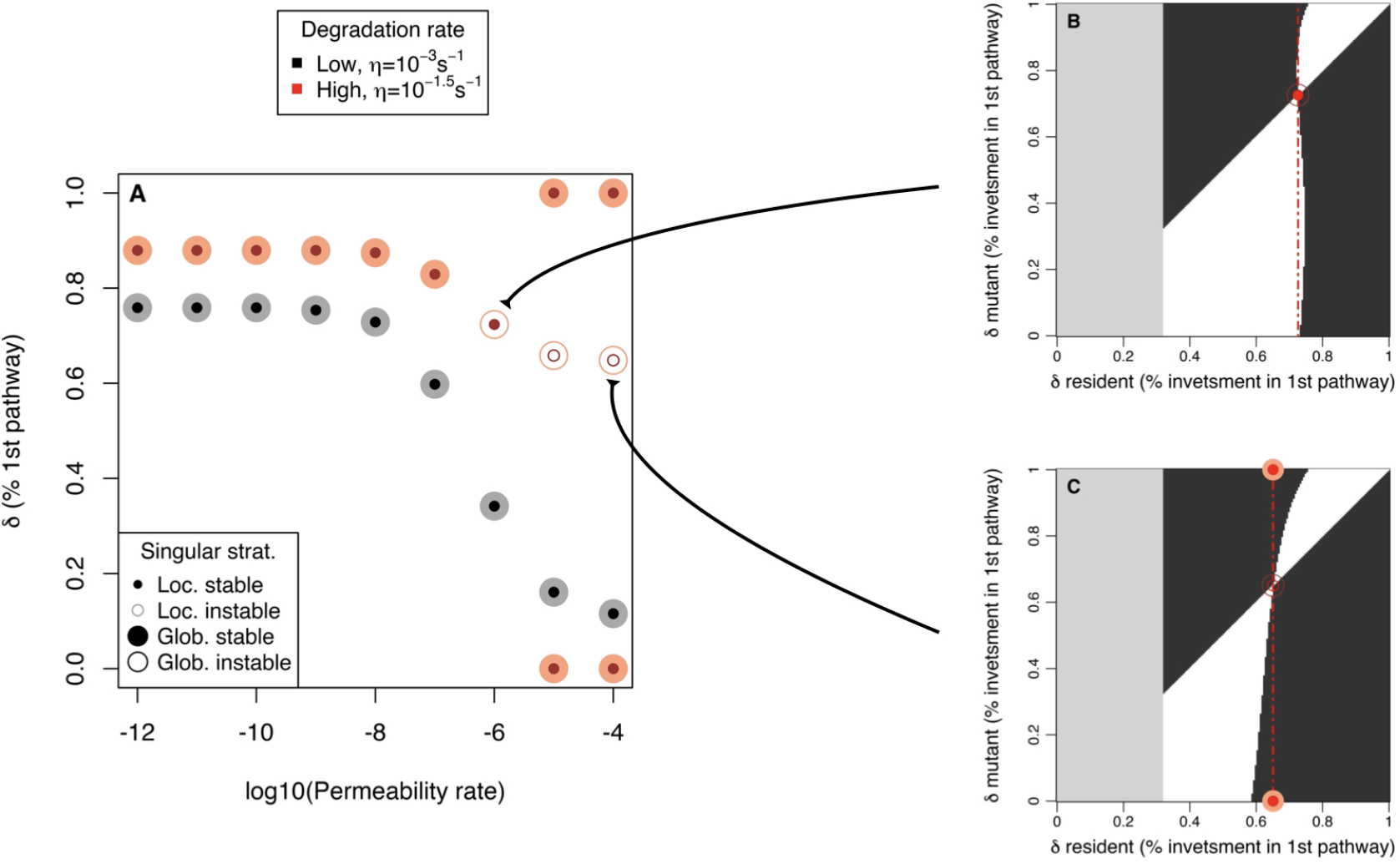
Permeability influences the evolution of strategies of enzyme allocations along a metabolic pathway, with the occurrence of cross-feeding at high permeability coefficients depending on the degradation rate. Two degradation rates are considered here (two other ones in Fig. S8 of SM) : low *η* = 10^−3^ s^−1^ in grey and moderate *η* = 10^−1.5^ s^−1^ in red. The enzymes catalysing reactions in either of two parts of the metabolic pathway, with their relative concentrations represented by *δ* (% 1st pathway), have kinetic parameters typical of those intervening in the carbon cycle (*k*_*f*_ = 10^6.25^ M^−1^. s^−1^ and *k*_cat_ = 10^2.25^ s^−1^); we also assume here that reactions in the second part *P*_2_ of the pathway produce 10 times more energy than reactions in the first part *P*_1_. For low permeability rates (below 10^−6^ dm.s^−1^), the evolutionarily expected strategy is always globally and locally stable – as in Figure 2 – and consists in investing more in the first part of the pathway as discussed in the text and described in further details through Fig. S5 and S15 (see the latter for the same enzyme efficiencies). Increasing permeability coefficients above 10^−7^ dm.s^−1^ results in a decrease in the investment in the first part of the pathway, *δ*, with different consequences for low and high degradation rates. At low degradation rates (in grey on panel A), lowering *δ, i*.*e*. increasing the investment in the second part, remains the most efficient strategy. At high degradation rates (in red on panel A), however, high permeability coefficients result in singular strategies that are both convergent (they evolve from any starting *δ*) and evolutionarily unstable (they can be invaded by mutants with close (locally unstable) or distant (globally unstable) allocation strategy *δ* – see panel C). This can lead to adaptive diversification, resulting in a stable coalition of strategies, that is, a population made of coexisting genotypes with different values of *δ*. The coalition can be determined (see SM - Figure S9); it is comprised of genotypes with *δ* = 1 (expressing only the first part of the pathway *P*_1_) and of genotypes with *δ* = 0 (expressing only the second part *P*_2_), corresponding to metabolic cross-feeding. Panel B represents a permeability *P* = 10^−6^ dm.s^−1^ for which the singular strategy is globally instable but locally stable, so that the evolutionary outcome should be contingent to the mutational landscape.

#### High permeability coefficients can promote cross-feeding

The overinvestment in the second part of pathway, *P*_2_, limits the loss of the leaky metabolite. As permeability increases, the investment in *P*_2_ must increase further, but this strategy becomes less efficient such that the external concentration of the metabolite rises. This, in turn, may give a selective advantage to genotypes that give up preventing the metabolite’s leakage by increasing their contribution to the first or second part of the pathway. From that point on, a new ecological niche may emerge, that can ultimately result in an evolutionary diversification between two genotypes. These situations can be identified by a specific type of pairwise invasibility plots where the singular strategy is convergent but evolutionarily unstable – as shown in Fig. 3-C for the case of a high degradation rate, the singular strategy at *δ* ≈ 0.65 can be invaded by nearby mutants – called a branching point. An adaptive diversification may occur at a branching point, requiring that we study the fate of mutants invading a pair of coexisting strategies, instead of a single one, through trait evolution plots (SM Figure S9).

Branching points indeed lead to an adaptive diversification as the diffusion of the intermediate metabolite increases above 10^−6^dm.s^−1^, with the two most extreme strategies, *δ* = 0 and *δ* = 1, evolving and forming a stable coalition (see SM - Text S3.1.2). This is characteristic of a complete metabolic cross-feeding, where a genotype transforms a nutrient into a metabolite released in the environment, and a second genotype feeds exclusively on that metabolite. For moderately high permeability rates – around *P* = 10^−6^*dm*. *s*^−1^ – the evolutionary outcome is more dubious as invasion is only possible by distant mutants; the result, in this case, will depend on the distribution of mutational effects and cannot be studied using classic adaptive dynamics methods.

It should be noted that the results in Figure 3 correspond to a metabolic pathway with unequal contributions of reactions to the energy needs of the cell (downstream reactions produce more than upstream ones, as observed on average in the carbon cycle). Equalling the energy contributions of all reactions often prevents the occurrence of cross-feeding. Yet, cross-feeding emergence was less sensitive to sub-pathway theoretical yields when we introduced the altogether realistic possibility of transporters and upstream enzymes co-expression, as exemplified on Figure S11. Reversibility and toxicity may also be involved in defining the critical point where cross-feeding coalitions can appear – see Section Text S3.3. Overall, the tipping point of *P* = 10^−6^ dm. s^−1^ seems robust across all internal selective environments, albeit not being a sufficient condition as it also depends on the specific combination of constraints associated with these environments.

More generally, the evolution of cross-feeding appears contingent on many factors. For instance, selection on the overall concentration of enzymes contributing to the pathway is important (*eg*. because it can reduce diffusive constraints), such that allowing higher concentrations precludes the occurrence of cross feeding. This is because increasing downstream concentrations, which may efficiently deal with metabolite diffusion, comes at a lower cost under these circumstances. This illustrates that the occurrence of cross-feeding may critically depend on the other tasks – and their contributions to fitness, which was not directly considered here, though the degradation rate may partly capture the behaviour – performed by cells and thereby on selection acting on their dedicated proteome, as the global proteome concentration may only vary to a small extent [40]. No less important should be the size of cells since smaller ones mechanically come with higher relative leakiness due to larger surface-to-volume ratios, which could in turn favor the occurrence of cross-feeding. Cells can nonetheless adapt their size to their content, at least to a certain extent [37], which may limit the costs incurred by an increase in concentration and prevent the occurrence of cross-feeding.

Finally, while in our model the efficiency of a reaction may only be changed through enzyme concentrations, kinetic parameters may also evolve and thus represent a relevant alternative in some specific parts of the metabolic network. The evolution of cross-feeding could thus also be contingent on the relative availability of mutations changing concentration *versus* kinetic parameters. Therefore, while this model includes much of the available information about enzyme kinetics and the selective constraints acting on the proteome, actually predicting how and when cross-feeding should evolve will require more efforts to better understand the building of global epistasis along metabolic networks and how critically this depends on the environment.

## Discussion

The fact that few metabolites – acetate and glycerol, noticeably – are more likely involved in the evolution of cross-feeding has been a conundrum for as long as experiments have revealed this phenomenon [25]. Here, we have put forward an explanation based on the necessity for a cell to optimise its proteome allocation, accounting for incurred costs and physical constraints like the differential permeability of a cell’s membrane to metabolites. Indeed, contrary to other metabolites, acetate is in constant chemical equilibrium with the highly diffusive acetic acid [42], and glycerol readily leaks towards the environment [40], which may predispose them to be involved in cross-feeding.

That metabolites rather upstream in a metabolic pathway – and therefore of potential use to generate more energy – will create a novel ecological niche when released is straightforward. But whether evolution will take this path when some genotypes have the potential to reduce the leak is worthy of a careful examination. In a number of cases that we have considered and that fall into the realistic range of parameters, proteome allocation will evolve in such a way that it prevents, or at least limits, the diffusion of the molecule. Only under some restricted conditions – that we have shown to coincide with the features of the two aforementioned metabolites – will cross-feeding evolve, characterised by a functional specialisation between a part of the population that transforms the nutrient into the diffusive metabolite, and another part that uses the metabolite as a carbon source, echoing work on digital evolution that similarly pointed to the possible contingency of cross-feeding [57, 58].

Beyond the existence of intrinsic cellular leakiness, one process that may facilitate the emergence of cross-feeding, or that may be co-opted to foster its efficacy when it is in place, is the cheap uptake and secretion of metabolites (*eg*. through facilitated transport) [42, 59, 60]. Even a slight leakage may for instance be enough to sustain a cross-feeder with high affinity transporters such as the one reported in *E*.*coli* for acetate [61]. This metabolic strategy, where cells actively give up a metabolite even though it still has the potential to bring fitness contributions, is often known as overflow metabolism [62, 63] and also frequently involves acetate. We did not consider the existence of a specific cost to the second part of the pathway, as has been documented in the past to explain overflow [64, 65]; nor did we account for other possible costs such as the existence of a localised toxicity [66] or the extra membrane occupancy involved in cellular respiration [67, 68]. These processes may promote the advent of cross-feeding, for they could bring extra fitness to an organism expressing one or the other part of the pathway.

From a broader ecological perspective, nutrients are hardly ever constantly renewed in nature; they are subject at least to some stochasticity, and most often than not occur through unpredictable periods of feast and famine. Insight into how proteome allocation may evolve in this context, and whether it should likely involve cross-feeding, can be gained from the existence of so-called diauxic shifts, where the phenotype can switch from the production of acetate to its consumption when the medium is enriched in this nutrient [59, 69]. In this case, it appears that the environment could be used to store intermediate metabolites, both increasing a genotype’s ability to uptake and use glucose when it is present, and its ability to await its renewal otherwise. Such plasticity in expressing different parts of a metabolic pathway should prevent, to some extent, the evolution of fixed specialists (as considered here) and further hinder the evolution of cross-feeding. We thus postulate that cross-feeding might only evolve in environments stable enough to prevent the evolution of plasticity.

Biology is actually teeming with interactions and emergent properties that complicate the big picture [70]. And, if functional simplicity may exist at the ecological scale [71], the ultimate underpinnings behind community assemblies are still blurred by the combination between a long and tumultuous evolutionary history and the wide prevalence of high order interactions, both within and between organisms, and in relation to the environment [57, 72, 73]. Addressing these community-level questions cannot bypass the existence of lower level features such as the epistatic relationships stemming from the joint effect of enzyme kinetic parameters [74] and their expression along pathways [75]; nor can we study metabolic evolution without a careful examination of their impact on the cellular micro-environment. We believe that the present study, by accounting for these interactions at a primordial ecological stage, will help explain how (some of) these communities appeared in the first place and how this may have then fuelled the scaffolding process underlying the building of microbial societies.

## Models

### Metabolic Model of fitness

Cell fitness results from the biomass and energy produced along a metabolic pathway (*eg*. ATP). The pathway is initiated by carrier proteins with *VT*_*m*_ = 1*mM/s* and *K*_*T*_ = 10*mM* (see SM Text S1 for details and justification), passively transporting nutrients inside cells and whose features are based on those for glucose in yeasts, as detailed elsewhere [34]. Nutrients are added (resp. eliminated) at a constant rate *α* (resp. *β*) in the external environment. The metabolic pathway is linear, comprising *N*_*r*_ (40 in the paper, but see SM for other values) reactions catalysed by enzymes whose levels of expression may evolve. The product of the *j*^*ieth*^ reaction is used as the substrate of the next one. These metabolites are constantly degraded at a rate *η* or fuelling toxicity *T* according to processes justified elsewhere [34] and also detailed at the beginning of the SM (introduction of section Text S1 and subsection Text S1.2). Each reaction (j) follows either reversible or irreversible Briggs Haldane kinetics [76]:

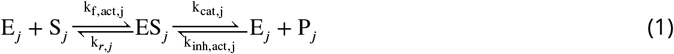

Irreversible reactions are modeled by setting *k*_*inh*_ = 0 – in this simple setting, the system can be solved for each pair of successive reactions since they do not feedback on upstream reactions, so that one has to solve *N* quadratic equations of the form:

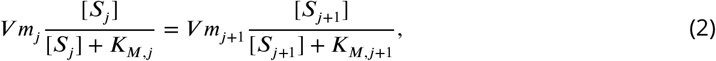

where *V*_*m*_ and *K*_*M*_ depends on microscopic parameters *k*_*f*_ and *k*_*cat*_, as well as enzyme concentrations while *η* is the degradation rate (leakiness is modelled in a similar manner than the degradation rate but depends on the gradient between internal and external concentrations so that it also impacts the environment through the latter). Otherwise, reversibility is spread equally between backward parameters *k*_*r*_ and *k*_*inh,act*_ (*eg*. if *K*_*rev*_ = 1*/*9, *k*_*r*_ = *k*_*cat*_*/*3 and *k*_*inh,act*_ = *k*_*f,act*_*/*3) and the system of *N* equations is solved using Broyden’s method [77]. The actual values for *k*_*inh,act*_ and *k*_*f,act*_ depend on the cellular constraints – see next subsection of Model.

Each reaction produces energy. This is a simplification as some reactions in the carbon cycle do not, unimportant as we consider the global expression of large portions of a pathway. We consider the case where contributions are equal along the pathway as well as other more realistic setups.

### Cellular constraints

Cell proteomes face two intrinsic constraints: (i) the cost of protein expression and (ii) the burden entailed by molecular crowding. We model (i) through a cost linear with protein expression, proportional to constant *c* [36]. We considered values of *c* such that the whole cytoplasmic proteome the enzymes in the pathway and other free enzymes – costs 5 to 50% of the whole cell budget. We model molecular crowding (ii) through a non-linear decrease of diffusion [38, 39] that changes the affinity constant *k*_f_ to *k*_f,act_ in the model with irreversible reactions, according to equation:

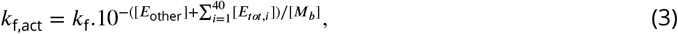

where [*E*_*tot,i*_] = [*E*_*i*_] + [*ES*_*i*_],[*M*_*b*_] = 3 · 10^−3^*M* represents the scaling factor for the effect of diffusion, while [*E*_other_] denotes the sum of the concentrations of other cytoplasmic proteins than the 40 under consideration. In the model with reversible reactions, *k*_*inh*_ is also affected by crowding, which is a conservative assumption as reverse reactions might be less sensitive to the diffusive process owing to the preexisting co-localisation between substrate molecules and enzymes.

Our model includes three processes involving the metabolites produced that select for the enhancement of enzyme activity, drawing a complex trade-off on the coexpression of enzymes: (1) metabolites can be lost, either because they are involved in parasitic reactions or because they are subject to targeted degradation [78], modelled through a linear degradation rate *η*; (2) metabolites can be toxic for the cell, for they engage in parasitic reactions, for instance through promiscuous interactions [79]; (3) highly reversible reactions within a pathway may also require efficient enzymes to maintain a high net flux [80]. We considered these three processes in various instances of our model, as described in SM Text S1; the results presented in the paper mainly comprise the action of a linear degradation rate, which provides a good qualitative understanding of how processes impacting the metabolites also impact selection on enzymes. Finally, the permeability of cell membrane to a given metabolite also acts as a constraint, which is introduced here by considering that one metabolite in the pathway diffuses passively at a rate *η*_*d*_ – on Figure 1, we show where this process occurs.

### Ecological equilibria

As to perform the adaptive dynamics analysis, we need to determine the ecological equilibrium set by a given genotype or coalition. To this aim, we first compute the value of the net flux (number of energy units per individual per unit of time) Φ_*net,N*_ for a given population size *N*. As long as this flux is lower (or higher) than an arbitrary value Φ_*net*_ = 10^−4^*M* · *s*^−1^, *N* is increased (or decreased) at the next iteration (by one unit when the flux gets close enough from this value). The algorithm is stopped, either when the difference between these fluxes is lower than 10^−6^Φ_*net*_, or when it oscillates between two neighbouring values (*N*_*eq*_ is then set to the average between these values). The ecological equilibrium, is defined by the nutrient and metabolite concentrations in the environment at this demographic equilibrium. Fitness values of strategies are then calculated in these conditions of equilibrium.

### Adaptive Dynamics of enzyme expression

We use Adaptive Dynamics [32, 43, 44] to model the evolution of enzyme expression along the metabolic pathway. This framework consider rare mutations, such that a resident “strategy” – corresponding to a given expression pattern – is assumed to have reached its demographic equilibrium before a mutant strategy appears in the population. At this demographic equilibrium, births compensate for deaths in the population, resulting in a concentration of nutrients specific of the resident (see previous section). The fitness of any mutant strategy is then determined for each resident equilibrium, which enables the drawing of Pairwise Invasibility Plots (PIPs) representing for each pair the ability of a rare mutant to invade the resident strategy, based on a comparison between the growth rate of the mutant and that of the resident. These plots are used to identify singular strategies and their properties, as defined in [43]. A particular type of singular strategies, branching points, may be indicative of a diversification in the population, which we further study by drawing areas of mutual invasibility (where both strategies invade when rare), before computing the ecological equilibrium for each coalition – composed of two resident strategies instead of one in that space. We then calculate the growth rate of mutants nearby each strategy in the coalition to identify coalitions that are stable (they cannot be invaded by any of the nearby mutants) and convergent (there exist evolutionary trajectories towards them), hence identifying evolutionarily expected communities after diversification has occurred [81, 82]. This latter process is summarised on trait evolution plots – TEPs – that we have shown in SM.

PIPs were generally drawn for 250 strategies, unless lower resolution was sufficient to capture the trend. In order to determine the optimal allocation, we set the total proteome content to its optimal value as determined without the influence of permeability. An individual whose strategy is to invest as much in the first part of the pathway than in the second part corresponds precisely to this case. Notice that, besides using a two step process, solving the systems to find nutrient and metabolite concentrations required to use R package ‘nleqslv’ with Broyden’s method [77].

### Settings for the models

A list of the basic settings can be found at the end of section Text S1.1 - SM. We varied them within their biological realistic ranges. This allowed us to identify key drivers of the diversification process that eventually result in cross-feeding, as discussed in the results section. The extensive analysis of parameters can be found in SM, especially in Texts S1 and S3.

## Supporting information

Supplementary materials - Proteome allocation and the evolution of metabolic cross-feeding

## Acknowledgment

We thank T.Lenormand for helpful discussions that led us to consider the evolution of cross-feeding. This preprint was created using the LaPreprint template (https://github.com/roaldarbol/lapreprint) by Mikkel Roald-Arbøl.

## Author contributions

FL, ER, FM and VD designed the research. FL and ER developed the models and analyzed the results. FL and ER wrote the first draft while FM and VD contributed further improvements.

