## Supplementary materials - Proteome allocation and the evolution of metabolic cross-feeding for "Proteome allocation and the evolution of metabolic cross-feeding"

#### Text S1 Total optimal content and cellular constraints

Numerous constraints may affect the total optimal content of cells. Dill et al. [2011] have shown that the amount of proteins should establish at an optimal intermediate level due to the deleterious impact an extra expression of proteins would have on diffusion. However, they did not account for the protein burden of producing these molecules. Their focus was besides on the total content without regard to the specific allocation of this content. This matters because a cell should be more prone to invest in a task when it entails large increases of fitness, and/or when these tasks are necessary to survive (meaning fitness is null in their absence). Here, we first assessed the effects of cellular constraints by considering a single pathway comprised of $N$ enzymes – see Figure S1.

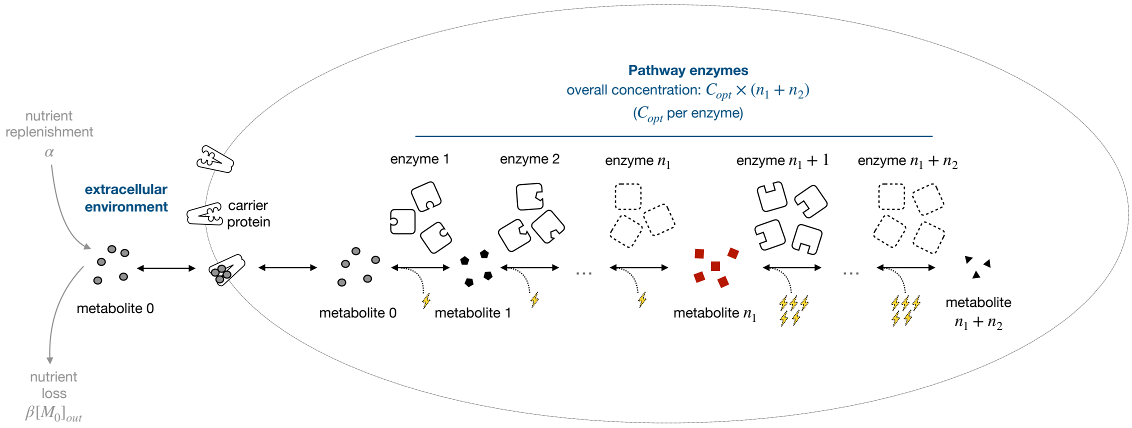


Figure S1: Overview of the model: the pathway is initiated by a carrier protein and comprised of N enzymes (*i.e.* corresponding to $n_1+n_2$ in the article). The extracellular dynamic of the nutrient is based on a constant replenishment-degradation process, where the loss is therefore proportional, through $\beta$, to the amount of nutrient. Within this pathway, each reaction follows Briggs Haldane kinetics (or its simpler Michaelis Menten form when reversibility is not accounted for) where enzyme efficiency is constant and studied as a parameter while reversibility varies depending on the subsection. These reactions each provide a fitness yield, shown above as lightning, proportional to the amount of product produced and to the specific yield of energy gain set for reactions – see section on overexpression. Fitness is simultaneously impeded by the cost of expression – production and crowding – and, depending on the subsection again, on the toxicity induced by the total concentration of metabolites.

We modelled a pathway initiated by a glucose carrier protein that facilitates diffusion, whose features correspond to average values – $V_{Tm} = 1mM/s$, $K_T = 10mM, \alpha = 1$ – for those reported in yeasts [Teusink et al., 1998, Maier et al., 2002]; notice that we do not report results of the influence of transporters since it had none on the processes we are interested in – it could have a significant influence in fluctuating environments, circumstances that are beyond the scope of this study. The chemical equation for facilitated transport can be approached by the following equation [ter Kuile and Cook, 1994, Bosdriesz et al., 2018]:

$$\frac{d[S_{\text{in}}]}{dt} = V_{Tm}.\frac{[S_{\text{out}}] - [S_{\text{in}}]}{K_T + ([S_{\text{out}}] + [S_{\text{in}}]) + \alpha.\frac{[S_{\text{out}}][S_{\text{in}}]}{K_T}} \tag{S1}$$

To match with central carbon metabolism, composed by glycolysis and the tricarboxylic acid cycle, the pathway modelled is comprised of $N = n_1 + n_2$ reactions, where $n_1$ and $n_2$ represent the respective number of reactions of two sub-pathways. Each reaction involves a specific enzyme and obeys Michaelis Menten kinetics, according to the following scheme (we relax the absence of reversibility later on, by using Briggs-Haldane equations [Briggs and Haldane, 1925]):

$$\mathrm{E_i} + \mathrm{S_i} \underset{\mathrm{k_r}}{\overset{\mathrm{k_{f,act}}}{\rightleftharpoons}} \mathrm{ES_i} \xrightarrow{\mathrm{k_{cat}}} \mathrm{E_i} + \mathrm{P_i}, \tag{S2}$$

where $k_{f,act}$ is the *in vivo* value of $k_f$ when accounting for the influence of crowding on diffusive processes. This influence is modelled by the following equation, already justified elsewhere [Labourel and Rajon, 2021]:

$$k_{f,act} = k_f \cdot 10^{-([E_{basal}] + \sum_{i=1}^{n_1+n_2} [E_{tot,i}])/[M_b]}, \tag{S3}$$

where $[E_{tot,i}] = [E_i] + [ES_i]$,$[M_b] = 3\ mM$ and represents the scaling factor for the effect of diffusion, while $E_{basal}$ – assumed to be constant – denotes the amount of protein allocated to other tasks than those of central carbon metabolism. Notice that compared with the previous reference [Labourel and Rajon, 2021], where we were mostly interested in setting a qualitatively realistic crowding limit, we have here refined this estimate to get as close as possible from physical findings [Blanco et al., 2018, Andrews, 2020] and from a realistic cellular protein fraction [Ellis, 2001, Dill et al., 2011]. Noticeably, the effect appears somewhat higher than previously, reflecting the need of an increase in the expression of cellular machineries such as ribosomes [Klumpp et al., 2013, Kafri et al., 2016] in order to produce more proteins.

Unless otherwise stated, $k_f$, $k_{cat}$ and $k_r$ are first set to values in line with (half an order of magnitude above) their median values estimated for central carbon metabolism [Bar-Even et al., 2011] – see next subsection. Remark that we test the sensitivity of the concentration ceiling to these values.

The selective pressure can be approximated by a linear degradation parameter $\eta_d$ competing with enzymes for their substrate (often denoted as a product since it also coincides with the product of the previous reaction), according to the following scheme:

$$\begin{array}{ccccc} \mathrm{P}_i + \mathrm{E}_{i+1} & \underset{k_r}{\overset{k_{f,act}}{\rightleftharpoons}} & \mathrm{E}_{i+1}\mathrm{P}_i & \xrightarrow{k_{\text{cat}}} & \mathrm{E}_{i+1} + \mathrm{P}_{i+1}, \\ \downarrow \eta_d & & & & \\ \mathrm{P}_{i,out} & & & & \end{array} \tag{S4}$$

where $P_i$ is the product of the previous enzymatic reaction and $E_{i+1}$ is the focal enzyme of the pathway (notice that the gap between indexes of $E$ and $P$ materialises because the first enzyme $E_1$ produces the first product $P_1$ from the substrate $S$), specialised at processing this product. As previously stated, the degradation rate $\eta_d$ that competes for metabolites applies to each substrate (including all intermediate products) in the pathway.

In the first section below, we test the influence of the selective pressure imposed by the degradation rate under different assumptions. As mentioned in the manuscript, our Adaptive Dynamics model relies on the dynamic competition between cells, and, more specifically, on the capability of mutants to invade resident strategies when these mutants are rare. The approach therefore account for the influence a resident strategy has on its environment.

#### Text S1.1 Influence of the degradation rate on optimal enzyme concentration

Because Adaptive Dynamics rely on the resident strategy having reached its ecological equilibrium, it is necessary to set an amount of net energy produced – the energy produced minus expenses entailed by protein production – that corresponds to this equilibrium whereby births exactly compensate for deaths. We set this amount to $\Phi_{eq} = 10^{-4}M.s^{-1}$, which leads to extracellular substrate concentrations close to the saturating constant of transporters. This parameter only influences the relative cost dedicated to the sustainment of the proteome (without any growth), and thence, the width of the viability area in the allocation space (as a higher sustainment cost decreases the energy that a cell can allocate to growth and reproduction).

Besides, there is also a need to set the size of cells – here, $r_c = 1\mu m$ corresponding to a volume of $V_c \approx 4.2\mu m^3$ (roughly that of *E.coli* cells) – and to consider a specific fraction of the environment – set to $V_{env} = 1000\mu m^3$ without cells (it is the volume of the environment that is free of cells). Notice that the size of cells matter when we study the influence of permeability inasmuch as passive diffusion depends on the SA:V ratio, which decreases when cells are bigger – this is discussed in the section about cross-feeding in the main document. On the contrary, the size of the environment does not matter: it only modifies the number of cells coinciding with the ecological equilibrium, which, in Adaptive Dynamics, is not involved in the outcome because genetic drift is ignored. Finally, the "chemostats" parameters are set such that the value of the enrichment rate equals that of the dilution rate, $\alpha = 10^{-3}\ Ms^{-1}$ and $\beta = 10^{-3}s^{-1}$ yielding a steady-state concentration in a cell-depleted medium of $[S_{out}]^* = 1M$. These coefficients are in line with estimates for diffusion coefficients of metabolites in solvant. Changing them may impact the equilibrium nutrient concentration or the speed at which nutrients are renewed in the environment. This may in turn influence the minimum flux that cells need to sustain and influence population viability. The set of parameters that are fixed at this stage is summed up in the following table:

| Parameters | $\Phi_{eq}(M^{-1}s^{-1})$ | $V_c(\mu m^3)$ | $V_{env}(\mu m^3)$ | $\alpha(Ms^{-1})$ | $\beta(s^{-1})$ |
|---|---|---|---|---|---|
| Values | $10^{-4}$ | 4.2 | 1000 | $10^{-3}$ | $10^{-3}$ |

Table S1: Set of constant parameters used to simulate competition

The system reaches an ecological equilibrium that needs be solved numerically through a two step process – see Model section of the article for details – such that it is not possible to determine the joint influence of the whole set of parameters. Instead, we varied them on a pairwise basis where the degradation rate is always the focal variable while other parameters but one (sometimes two) are fixed. When not explicitly mentioned, these parameters are set to their default values in the following table, where the pathway yield represents the total amount of energy provided by the pathway for the processing of one glucose molecule (the yield is spread equally among the reactions, unless otherwise stated):

| Parameters | $k_f(M^{-1}s^{-1})$ | $k_{cat}(s^{-1})$ | $c_{exp}$ | $E_{basal}(M)$ | Pathway yield |
|---|---|---|---|---|---|
| Default values | $10^{6.5}$ | $10^{2.5}$ | $10^{-2.5}$ | $5.5 \cdot 10^{-3}$ | $1G : 10E$ |
| Range of variation | $10^6 - 10^7$ | $10^2 - 10^3$ | $10^{-3} - 10^{-2}$ | $5 - 6 \cdot 10^{-3}$ | $1G : 5E - 20E$ |

Table S2: Set of basic settings along with the range of variations used to determine their influence in isolation

Notice also that in this first subsection, $k_r$ is always set to equal $k_{cat}$, an assumption relaxed when reaction reversibility is introduced, according to modalities described in the relevant section.

##### Text S1.1.1 Influence of enzyme kinetic parameters

Adaptive enzyme concentration along the pathway

To evaluate how kinetic parameters impact the optimal cell content, we modeled the evolution of enzymes along a metabolic pathway for three different values of enzyme efficiency in line with the literature [Bar-Even et al., 2011, Labourel and Rajon, 2021], varying them by half an order of magnitude around (and slightly above, so as to be in conservative cases for which the content is facing a rather moderate selective constraint) their interspecific median values for central carbon metabolism.

At this stage, three selective pressures are considered:

i) the degradation rate competing with enzymes for metabolites;

ii) the cost of protein production, be it due to production or because of molecular crowding;

iii) the level of flux.

We consider all enzyme concentrations belonging to a pathway to be identical, and show that cells consistently evolve to invest more than 15% of their proteome content to the pathway – see Figure S2-A below, where even cells facing low degradation rates invest at least 15% of their content.

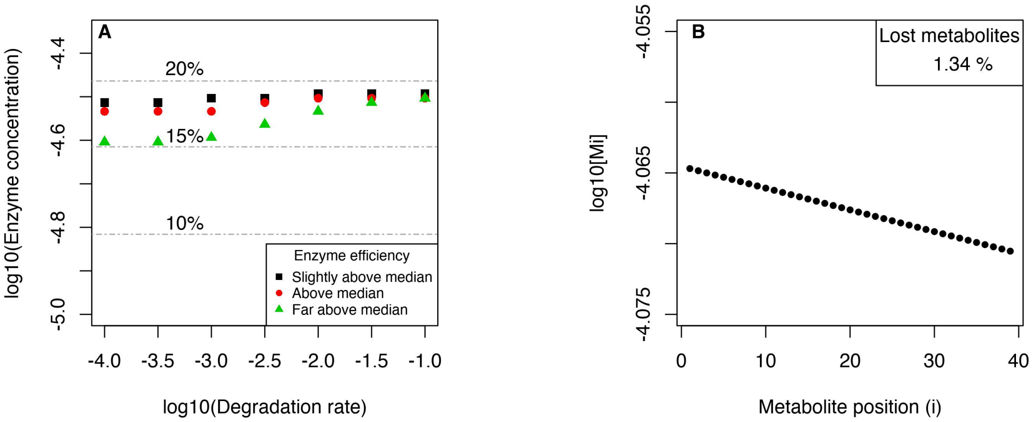


Figure S2: Influence of kinetic parameters on the optimal content of cells. The optimal concentration that each enzyme needs reach is represented for slightly above median (black squares) – $k_f = 10^6 M^{-1} \cdot s^{-1}, k_{cat} = 10^2 s^{-1}$, above median (red circles) – $k_f = 10^{6.5} M^{-1} \cdot s^{-1}, k_{cat} = 10^{2.5} s^{-1}$ – and far above median (green triangles) – $k_f = 10^7 M^{-1} \cdot s^{-1}, k_{cat} = 10^3 s^{-1}$ – enzyme efficiencies. In A, as the degradation rate increases, the selective pressure on the content gets a little higher, and thereby the optimal enzyme concentration, finally hitting a ceiling. Notoriously, the increase of the content eventually cannot overcome a given ceiling in any case – when accounting for 20% of the proteome – where the effect of hindered diffusion always overcomes the extra gain of activity. The increase of the adaptive expression occurring with the degradation rate is limited because the degradation rate has a combined antagonistic effect: it enhances the pressure to limit metabolite loss but it also induces the latter so that more downstream reactions experience a lower selective pressure. The loss of metabolites along the pathway is log-linear as exemplified in B for the lowest degradation rate $\eta = 10^{-4} s^{-1}$, and an enzymatic concentration of $10^{-4.5} M$ for each reaction.

Increasing the degradation rate of metabolites enhances the selective pressure for enzyme efficiency, which can only improve here through an increase in enzyme concentration. Overall, the adaptive enzyme content reaches a ceiling close to 20% of the total proteome content (see Figure S2-A); at this point, molecular crowding becomes a major constraint preventing further improvement. The increase in proteome allocation is limited because of the antagonistic effect of the degradation rate, that on the one hand enhances the selective pressure to avoid metabolite losses but on the other hand also reduces the selective pressure acting on downstream enzymes because of the resulting lower flux of metabolites, as we illustrate below.

##### Influence of the degradation rate on the concentrations of metabolites along the pathway

The loss of metabolites along the pathway obeys a log-linear effect (see Figure S2-B) that may eventually sum to high overall values. Here we quantify this loss as one minus the ratio of the last metabolite concentration to the nutrient concentration (Table S3).

| $[E_1]_{tot}$ / $\log_{10}(\eta)$ | $10^{-4.5} M$ |
|---|---|
| – 4 | 1.34% |
| – 3.5 | 4.16% |
| – 3 | 12.56% |
| – 2.5 | 34.47% |
| – 2 | 73.3% |
| – 1.5 | 98.2% |
| – 1 | 100% |

Table S3: Metabolite losses for a wide range of degradation rates, $\eta$.

Up to $\eta < 10^{-2} s^{-1}$, the situation may coincide with realistic losses of metabolites [Park et al., 2016]. Higher degradation rates ($\eta \geq 10^{-2} s^{-1}$ ; orange and red values in Table S3 appear unrealistic under this model, but may nonetheless describe appropriately the selective pressure on enzymes in a more realistic context (for instance, reversible reactions or branching in the metabolic network; Labourel and Rajon 2021).

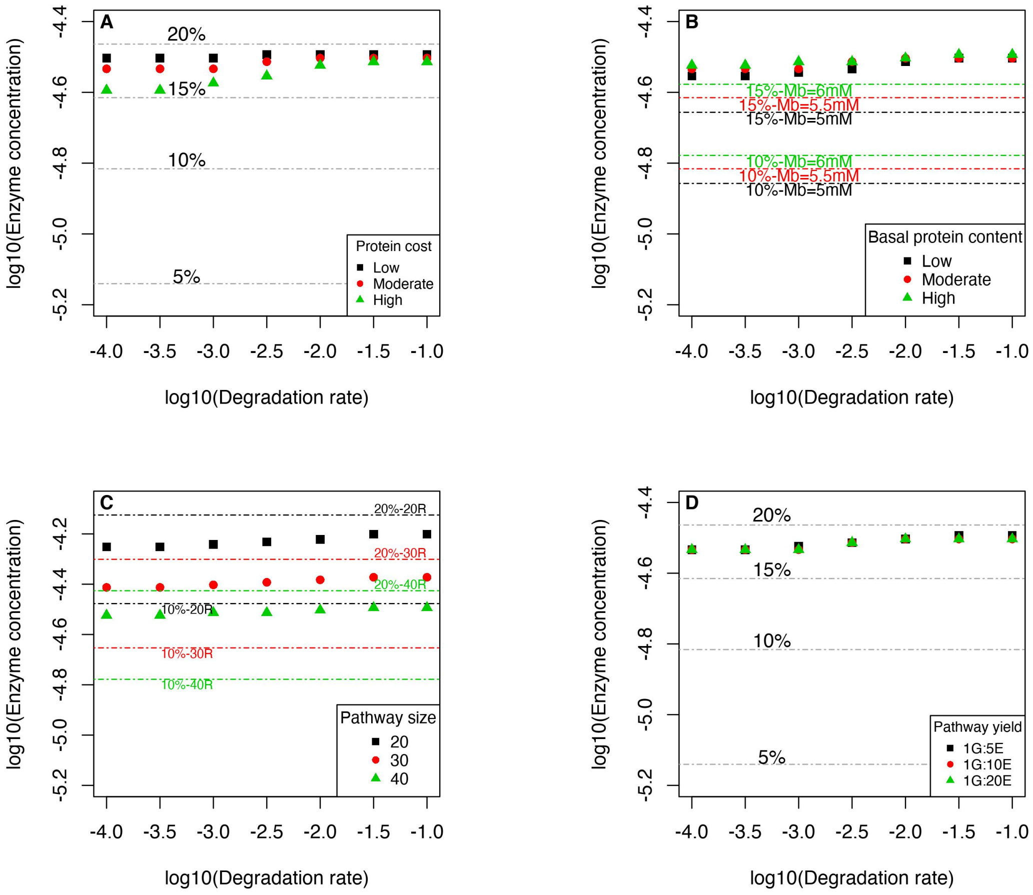


Figure S3: Influence of cellular parameters on the optimal proteome content has proven to be minor when all enzymes covary together. (A): linear protein cost, where a low cost means that intracellular proteins approximately accounts for 5% of the total budget of cells while a high one approximately accounts for 50% of the same budget. (B): basal proteome concentration [$E_{basal}$] (proteome fraction not dedicated to energy metabolism like the central carbon metabolism, see Equation for $k_{f,act}$ above) – the total fraction of the proteome also depends on the background concentration, which explains why distinct lines are drawn. (C): pathway size ranges from 20 to 40 reactions and, again, the total fraction of the proteome depends on the focal variable, explaining the distinct lines depicting the relative investment represented by the focal pathway. (D): pathway yield varies from 5 to 20 "fitness" molecule(s) per glucose molecule (for the pathway as a whole, meaning that the yield of one reaction is this amount divided by the size of the pathway). Notice, that an energy molecule can be 1 ATP, for instance. In each of these cases, the total fraction of the proteome dedicated to energy metabolism cannot exceed an amount between 15% to 20%.

##### Text S1.1.2 Influence of other cellular parameters

As to study the influence of other parameters, we then set $k_f = 10^{6.5}M^{-1}s^{-1}$, $k_{cat} = 10^{2.5}s^{-1}$ and $k_r = k_{cat}$, approximately half an order of magnitude higher than median estimates of Bar-Even et al. [2011]. As previously, the concentration of the upstream enzyme is considered to covary with the other ones. In this subsection, we tested the influence of the major cellular and metabolic parameters and demonstrated that the main influence is still set, as expected following the previous results obtained, by the degradation rate the transporter protein (see Figure S3).

##### Text S1.1.3 Influence of environment parameters

The replenishment of the environment results from the flux parameter $\alpha$ while the degradation results from the rate parameter $\beta$. They had no impact on the optimal content, only changing the demographic equilibrium (with a lower $\alpha$, the steady-state population diminishes and may even vanish, even if the steady-state concentration in the environment is high). We do not report these results but the parameters are easily handled in the dedicated scripts.

#### Text S1.2 Influence of more realistic set of constraints on the optimal content

We determined the effect of toxicity using a non-linear effect – as in Chou et al. [2014] – affecting fitness according to the following equation:

$$f = (\Phi - cost \cdot \sum_{i=1}^{40}[E_{tot,i}]) \times \frac{T}{T + \sum_{i=1}^{40}[M_i]},$$

where $T$ is a toxicity constant, which, when reached by the total metabolite content, cut fitness by half.

For example, a toxicity constant set to $10^{-1}M$ means that if the sum of all 40 metabolites involved in the pathway equals this amount, then fitness is half what it would have been without this constraint. Toxicity has a similar qualitative effect than that of degradation – see Figure S4-A, where T varies from $10^{-2}M$ to $1M$: independently of any other factor, a high toxicity pushes the selective pressure much similarly than the degradation rate does. The maximal optimal content – coinciding with approximately 15-20% of the pathway dedicated to the energy metabolism – does not differ from the ones yielded by other parameters.

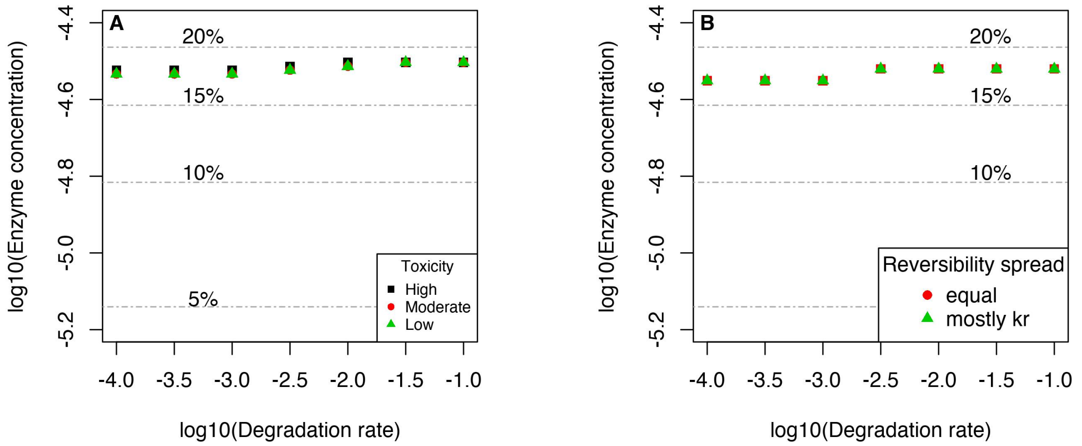


Figure S4: Influence of metabolite toxicity (A) and reversibility (B, combined with a basal toxicity) on the optimal content. Toxicity T – High: $T = 10^{-2}M$; Moderate: $T = 10^{-1}M$; Low: $T = 1M$ – increases the selective pressure acting on enzyme concentration in A, so that even with a low degradation rate, the adaptive cell content is close to its adaptive value with a high degradation rate. Assuming a moderate toxicity – $T = 1M$ – in (B) that combines with a realistic level for reaction reversibility, the selective pressure is again less dependent on the degradation rate, and changes in where the reversibility is focused – "equal": $k_r = k_{cat}/3$, $k_{inh} = k_f/3$; "mostly $k_r$" :$k_r = k_{cat}$, $k_{inh} = k_f/9$ – only impacts the viability of cells (when relying mostly on $k_{inh}$, cells are no longer viable - results not displayed here).

In parallel, reversible reactions obey the following scheme (where (i) denotes the $i^{eth}$ reaction):

$$\mathrm{S}_i + \mathrm{E}_i \underset{k_{r,i}}{\overset{k_{f,i}}{\rightleftharpoons}} \mathrm{E}_i S \underset{k_{inh,i}}{\overset{k_{\mathrm{cat},i}}{\rightleftharpoons}} \mathrm{E}_i + \mathrm{P}_i \tag{S5}$$

Reversibility was therefore studied by considering that it can affect either $k_r$, $k_{inh}$ or both. The level of reversibility was set to a specific value ($K_{rev} = 1/K_{eq} = [S]_{eq}/[P]_{eq} = 1/9$), which is the geometric mean of that for reactions involved in the central carbon metabolism and whose values have been summarised in Li et al. [2011]. As reversibility can be spread between two parameters, it is necessary to see how the flux reacts to this intrinsic process under the cellular constraints. Because $k_{inh}$ is an association constant, it is subject to the effect of crowding on diffusion much like $k_f$, according to equation (S3). Notice that we simplify the impact of this process of reversibility by assuming it does not evolve, even though it was shown that organisms should in principle optimise the energy profile determining how it is spread [Heinrich et al., 1991, Klipp and Heinrich, 1994].

If the reversibility impedes mostly the parameter $k_{inh}$, there is no degradation rate susceptible to produce a flux high enough to compensate for the need to sustain its pool of proteins. Therefore, we report results only for the two other cases where reversibility acts mostly on $k_{inh}$, or that for which it is equally spread between both parameters ($k_{cat} = k_r/3$ and $k_{inh} = k_f/3$). The effect of reversibility also increases the selective pressure, as shown in S4-B. In both cases, there is still a little room for extra protein expression when degradation rates are low. We do not report the influence of reversibility solely, for it proved to be similar. Last but not least, we also checked that the combined effect of reversibility and toxicity on enzymatic demand has a large effect (pushing towards high enzyme concentration approaching 20%) even when the expression of the first enzyme is set so that the high demand is not only the result of the relevance of a high upstream investment, but more generally that of the propagation of selective pressure along the pathway as theorised recently in Kryazhimskiy [2021].

### Text S2 Overexpression of upstream enzymes

In this section, we investigate why cells preferentially express upstream enzymes. We first study an irreversible pathway as to avoid a fuzzy picture resulting from too many parameters, before extending the framework to include other mechanistic constraints. The conclusion taht enzymes upstream are consistently over-expressed is robust to these changes.

#### Text S2.1 Case of an irreversible pathway

To determine how cells should allocate their proteome along a metabolic pathway, we consider that the pathway is comprised of two subpathways of respective sizes $n_1$ and $n_2$, and enzyme expression may differ between (but not within) each subpathway. The total available proteome expression for the whole pathway is set to its adaptive value $[E]^*$ obtained when reactions are assumed to be identical all along it – see previous sections for more details. The pathway can be described as follows:

$$\mathrm{S}_{out} \xrightarrow{T} \mathrm{S}_{in} \xrightarrow{E_0} \mathrm{P}_{1,1} \xrightarrow{E_{1,1}} \cdots \xrightarrow{E_{1,n_1-1}} \mathrm{P}_{1,n_1} \xrightarrow{E_{2,1}} \mathrm{P}_{2,1} \xrightarrow{E_{2,2}} \cdots \xrightarrow{E_{2,n_2}} \mathrm{P}_{2,n_2} \tag{S6}$$

In this chemical equation, transporters $T$ drive the uptake of substrate $S$ that is then transformed into product $P_{1,1}$ by the enzyme $E_0$. $(n_1 - 1)$ identical irreversible reactions belong to the upstream subpathway, along which enzymes of the same concentration $[E_1]$ transform $P_{1,1}$ into $P_{1,n_1}$. The same process occurs along the second subpathway, with $(n_2)$ enzymes of similar concentration $[E_2]$. In the model described in the main text, the first enzyme has the same, evolvable concentration as all the upstream enzymes (results shown in (figure S5-B for this case). But this enzyme has a specific role in the irreversible model: it secures nutrients that would otherwise be lost due to the reversible actin of the transporter, such that this particular selection pressure may drive the evolution of the concentration of its subpathway. In order to separate this from other selection pressures, we also considered a case where the most upstream enzyme has an independent concentration (Fig. S5-B). Remark that the number of evolvable upstream enzyme concentrations is thus either $(n_1 - 1)$ (A) or $(n_1)$ (B). From now on, we consider $n_2 = n_1$.

##### Text S2.1.1 Adaptive Dynamics outcomes

Irreversible subpathways always favour an overexpression of upstream enzymes. This conclusion is trivial when degradation rates are high – see (A) and (B) on Figure S5 – for the mere reason that there are fewer metabolites to process in the second subpathway due to upstream degradation – see Figure S2 for the illustration of metabolite losses. Nearly as predictable is the influence of a distortion in the supply of energy provided by reactions: when downstream reactions provides a higher pay-off, downstream expression increases. Yet, and even when the pay-off should lead to prioritise downstream reactions – see (A) on S5 for low degradation rates, where the investment in the first pathway is always more than half – it is always adaptive to allocate more to upstream ones: this is because within an irreversible pathway, upstream enzymes are involved directly in the fitness provided by their reactions and indirectly in that provided by the following ones, and not the opposite – see next subsection for a more detailed analysis of this effect. One final remark is that when the first enzyme is constrained to the same level than downstream ones, the adaptive investment is also shifted towards upstream reactions – see low degradation rates on Figure S5-B. This is because the first enzyme competes with transporters to process its substrate owing to the reversible nature of the uptake process, which enhances the selective pressure it experiences [ter Kuile and Cook, 1994, Labourel and Rajon, 2021] in a context where other enzymes are almost free of constraints. However, accounting for reversibility and toxicity of intracellular reactions removes the artificial specificity of the first enzyme (and transporters) when degradation rates are very low and it tends again to show that what matters is mostly about the balance between the contributions (both direct and indirect) of each subpathway – see the case of an irreversible pathway below.

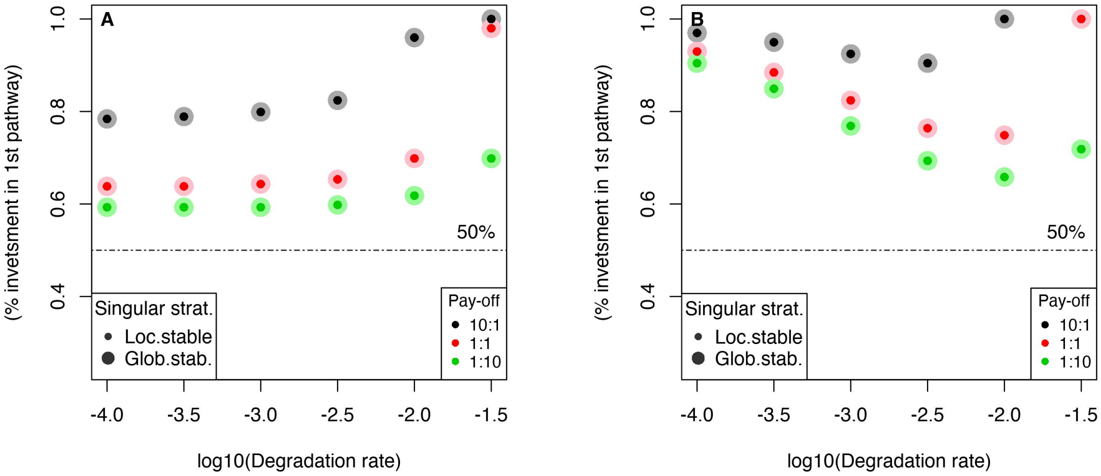


Figure S5: Selective constraints along an irreversible metabolic pathway push cells to focus on upstream reactions, no matter how energy gains are spread along the pathway – 10:1 as pay-off means that each reaction of the first subpathway provides 10 times the energy provided by each of the downstream reactions, and the other way around for 1:10. In (A), the influence of transporters is set aside by assuming a non-evolvable first enzyme concentration. Self-evidently, the degradation rate enhances the relevance to prioritise the first subpathway, because metabolites are progressively lost so that fewer metabolites can be processed downstream. More interestingly, even when the degradation rate is so low that very few metabolites are lost – see low degradation rates such as $\eta = 10^{-4}s^{-1}$ – it remains relevant to focus on the first subpathway (see text and next subsection for more details on this phenomenon). In (B), besides the two phenomenon described in (A), the impact on the transporter is shown to complexify the picture without changing the qualitative conclusion of an adaptive overexpression in upstream reactions: with low degradation rates, the transporter becomes the main constraint and because the selective pressure is low elsewhere in the pathway, it favors a very high overexpression of upstream reactions that tends to get lower with intermediate degradation rates because the seective pressure is homogenised along the pathway. Eventually, the loss of metabolites becomes so high that cells again should waive most if not any downstream investment.

##### Text S2.1.2 Differential allocation between subparts of pathways: a toy model

In this section, we introduce a toy model designed to unravel the intricacies behind the optimal allocation strategy along an irreversible pathway. Instead of considering a long pathway, we focus on a pathway made up of two consecutive reactions that contribute to fitness, where the flux prior to the first reaction is denoted by $\Phi_0$. Based on insights from the flux control theory [Kacser and Burns, 1973, Heinrich and Rapoport, 1974] and a more recent mechanistic approach [Labourel and Rajon, 2021] partly relaxing the need for unsaturated reactions, the flux sustained by enzymes of reaction (i) can be written as:

$$\Phi_i = \Phi_{i-1}\frac{[E_i]}{K + [E_i]}, \tag{S7}$$

where $[E_i]$ denotes the total concentration of enzyme (i) and $K$ represents a phenomenological saturation parameter acting on enzymes and involving different constraints emerging within a pathway [Hartl et al., 1985, Kaltenbach and Tokuriki, 2014].

With two reactions in the pathway, the system can be summarised as follows:

$$\begin{cases} \Phi_1 = \Phi_0\dfrac{[E_1]}{K_1 + [E_1]} & \text{(S8a)} \\ \Phi_2 = \Phi_1\dfrac{[E_2]}{K_2 + [E_2]} & \text{(S8b)} \\ W = \Phi_1 + \Phi_2 - c\cdot([E_1] + [E_2]), & \text{(S8c)} \end{cases}$$

with $c$ representing the cost of protein production and $\Phi_1$, $\Phi_2$ the fluxes that directly (and equally) contribute to fitness $W$.

Assuming that the parameter $K$ is identical for both reactions, fitness can therefore be written as:

$$W = \Phi_0 \cdot (\frac{[E_1]}{[E_1]+K})(1+\frac{[E_2]}{[E_2]+K}) - c \cdot ([E_1]+[E_2])$$

Note that what generates the flux $\Phi_0$ does not matter for our purpose, though it may be seen as the flux produced by carrier proteins translocating a specific nutrient.

Because the total enzymatic concentration $[E_{tot}]$ is constrained, let us assume that it is set to its optimal value. Thence, the optimal allocation stemming from such a system is reached when the extra fitness gained by increasing either one of the concentration equals that obtained through the other one. Indeed, at the point where any increase of the total concentration does not entail any extra fitness, this concentration has to be spread between pathways and if it promotes fitness to increase the allocation in one pathway, it implies that there is a corollary interest to decrease the allocation to the other one, as shown when writing the two corresponding equations:

$$\begin{cases} E_2 = E_{tot} - E_1 \Leftrightarrow \dfrac{\partial E_2}{\partial E_1} = -1 & \text{(S9)} \\ \dfrac{dW}{dE_1} = \dfrac{\partial W}{\partial E_1} + \dfrac{\partial W}{\partial E_2}\dfrac{\partial E_2}{\partial E_1} = 0 & \text{(S10)} \end{cases}$$

As a consequence, the condition can be written as:

**Condition 1** $\frac{\partial W}{\partial [E_1]} = \frac{\partial W}{\partial [E_2]}$

This condition straightforwardly requires the following quadratic equation to hold:

$$([E_1])^2 + K[E_1] - (2[E_2]+K)([E_2]+K) = 0,$$

which can be rewritten as:

$$[E_1] = \frac{-K + (K^2 + 4(2[E_2]+K)([E_2]+K))^{1/2}}{2} \tag{S11}$$

Finally, one can distinguish this optimal allocation depending on the level of saturation of the reactions:

$$\begin{cases} [E_1] \approx \sqrt{2}[E_2], \text{ if } [E_2] \gg K & \text{(S12a)} \\ [E_1] \approx 2[E_2], \text{ if } [E_2] \longrightarrow K & \text{(S12b)} \\ [E_1] \gg [E_2] \text{ and thus } [E_1] \text{ is mostly independent from } [E_2], \text{ if } K \gg [E_2]. & \text{(S12c)} \end{cases}$$

Denoting $\delta = \frac{[E_1]}{[E_{tot}]}$ the fraction of the proteome dedicated to the first reaction as in the main document and following a similar optimisation approach yields that the minimum adaptive overexpression $\delta^*$ is given by $\delta^*_{min} = 2 - \sqrt{2} \approx 0.585$ and coincides with saturated pathways, *i.e.* $[E_1] >> K$ and $[E_2] >> K$. Replacing $[E_1]$ and $[E_2]$ in $W([E_1],[E_2])$ with $\delta$ and $E_{tot}$, one can show that:

$$\delta^* = \frac{2(\theta_s+1) - \sqrt{(2\theta_s+3)(\theta_s+1)}}{\theta_s}, \tag{S13}$$

where $\theta_s = \frac{E_{tot}}{K}$ represents the level of saturation – see Figure S6 – for the shape of the optimal allocation for the toy model.

In brief, it means that a cell comprised of irreversible pathways should consistently allocate more to upstream reactions because they contribute more to fitness, a phenomenon that has a maximum impact when reactions stand far from saturation. The limit obtained with this model seems close to the floor value in Fig. S5, which we assume is fortuitous due to the distinct nature of the two models. In particular, the model presented in this section does not capture realistically reactions approaching saturation [Bagheri-Chaichian et al., 2003] – but see [Yi and Dean, 2019] and [Labourel and Rajon, 2021] for other approaches showing similar saturating effects of enzyme concentrations.

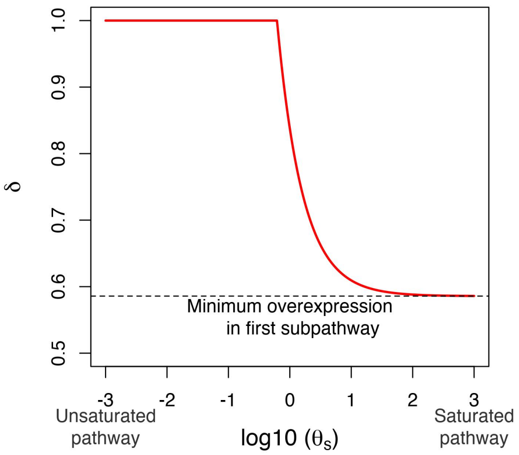


Figure S6: (A) shows the optimal strategy for different levels of pathway saturation $\theta_S$ (on a log-scale). Apart from a small area where reactions are close to the half saturation constant, the optimal allocation is either to prioritise fully the upstream reaction – case of unsaturated pathways – or to allocate slightly more to them – case of saturated pathways. This latter case corresponds to the minimum overexpression whose value is 0.585 – see text for details.

#### Text S2.2 Case of a reversible pathway

In this subsection, we report results obtained when considering a more realistic set of constraints combining both reversibility, toxicity – see Text S1.2 for details about how we model these processes – and degradation. Because what matters when reversibility enters into play is both the way reactions contribute to fitness and constraints set by the initiating transporter (whose reversibility rate is close to 1 in the case studied), the adaptive allocation still favors the expression in the upstream sub-pathway, despite the pressure also set by toxicity – see (A) and (B) on Figure S7. Notice that with reversibility, downstream reactions can contribute indirectly to the net upstream flux by limiting return fluxes. The relative weight of indirect contributions thus depend on the precise level of reversibility. When reversibility is below 1, metabolites are more quickly pushed downwards by upstream reactions than they are pulled upwards by downstream ones, which means that upstream reactions have a larger indirect influence on fitness than the opposite. Because reversibility is spread all along the pathway, the adaptive allocation quickly switches when $K_{eq} = 1$ to an equal allocation between downstream and upstream enzymes, while a higher reversibility would lead to the need of enhancing the expression of downstream enzymes, as indirect contributions of downstream reactions would exceed that of upstream ones. Notice that the (geometric) average reversibility of the central carbon metabolism is approximately $10^{-1}$ [Li et al., 2011] leading to a rather high departure from the equal spread. Assuming such a reversibility and different payoffs for each subpathway does not change much the optimum allocation, but a high generic – equally spread between each reaction – toxicity tends to homogenise how cells should allocate their proteome as to avoid that some metabolite concentrations go beyond a given level. It remains an open question beyond the scope of this study to determine how more heterogeneous (and realistic) constraints influence more specifically the allocation process.

### Text S3 Membrane permeability and optimal allocation between pathways

Because fitness contributions add up along the pathway, we have shown that it may be relevant for an organism to favour certain of its reactions over others. Up until there, the selective pressures faced by enzymes were identical. Yet, the situation may turn out very differently, for instance if a metabolite is susceptible to be released in the environment, either passively through simple diffusion or actively through excretion machineries. In this section, we evaluated the impact on optimal metabolic strategies that membrane permeability may beget.

#### Text S3.1 Model case – complementary results and plots

Unless otherwise stated, the total enzymatic content available for the focal pathway is set at its optimal value when no metabolite is subject to permeability. We examine this assumption in subsection Text S3.1.3 and show that it should not influence outcomes but marginally.

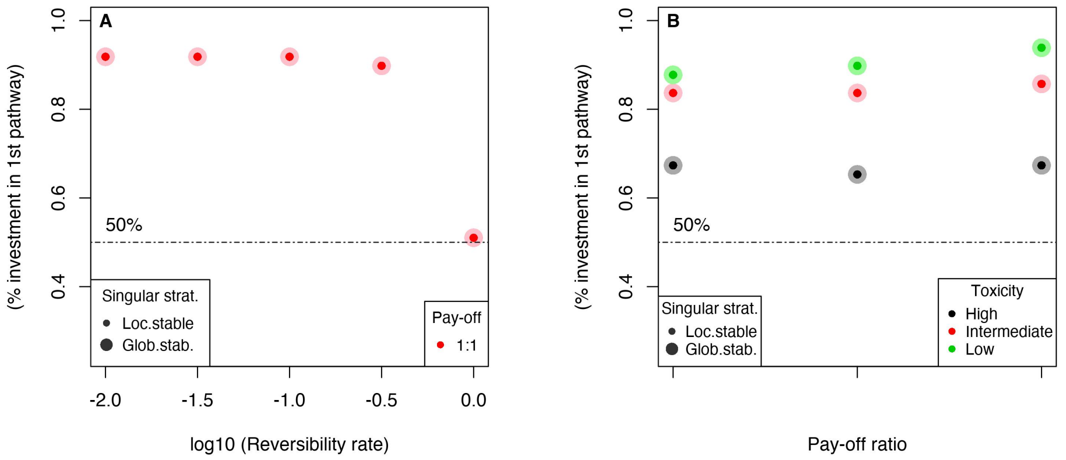


Figure S7: Influence of reversibility and toxicity on the adaptive allocation when degradation is negligible ($\eta_d = 10^{-4}s^{-1}$). In (A), the influence of reaction reversibility is shown for an equal yield of each subpathway when a low toxicity ($T = 1M$) is considered: as long as reversibility is below 1, the first subpathway should be favoured. In (B), reversibility is set to its geometric mean for enzymes involved in central carbon metabolism: no matter the toxicity level, the pay-off ratio between sub-pathways yields is proven to be of little influence. Although the way toxicity is modelled does have an effect because it pushes pathways to homogenise their metabolite levels, this does not undermine the idea that cells should prioritise upstream reactions.

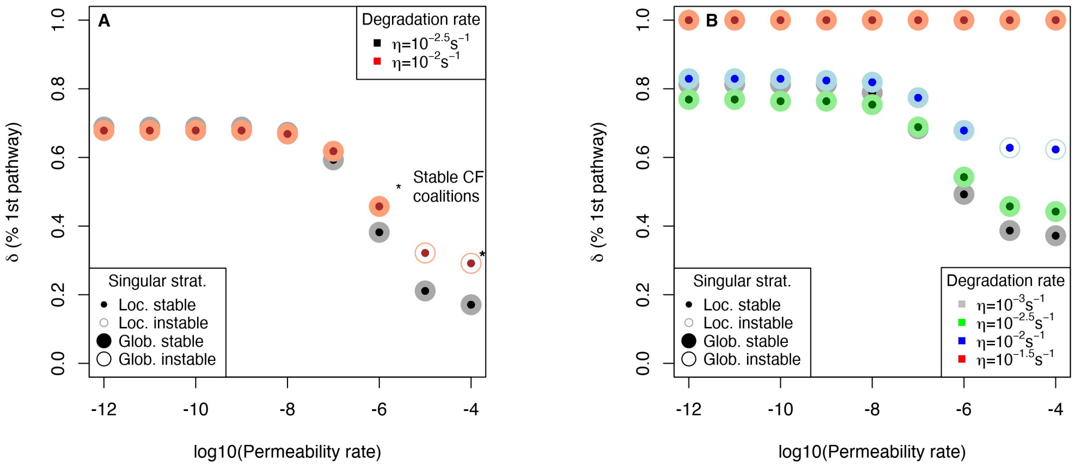


Figure S8: In (A), the plot shows the strategy that evolved when considering different degradation rates ($\eta = 10^{-2.5}s^{-1}$ and $\eta = 10^{-2}s^{-1}$), a 1:10 pay-off spread between sub-pathways and enzymes moderately efficient (with regards to values found for the central carbon metabolism), for different permeability levels of the membrane – permeability only concerns one metabolite, in the middle of the pathway. To cope with this phenomenon and avoid the cost of leakiness, cells allocate more to the second part of the pathway in either cases, starting from an overexpression of upstream enzymes ($\delta > 0.5$) with low permeability to the opposite allocation when it exceeds $P = 10^{-6}dm \cdot s^{-1}$ ($\delta < 0.5$). Singular strategies are convergent and stable, except for $\eta = 10^{-2}s^{-1}$ and $P > 10^{-6}dm \cdot s^{-1}$, where an ecological niche tends to emerge and lead to cross-feeding coalitions with the highest value of $P$, denoted with the star. With $P = 10^{-5}s^{-1}$, both outcomes are possible and depend, amongst other things, on the distribution of mutation effects. In (B), we consider a 1:1 pay-off that narrows down the possibility for mutual invasibility, and restricts possible coalitions to generalist/specialist of the initial resource, which in turn implies that when the degradation rate is high enough to always favour these latter, no coexistence can emerge (see $\eta = 10^{-1.5}s^{-1}$). Therefore, these conditions do not enable CF interactions.

##### Text S3.1.1 Outcomes with degradation only along the pathway

We first report results – see Fig. S8-A – obtained for the model case studied in the main document. It is based on an irreversible pathway initiated by a reversible transporter and whose metabolites are subject to degradation. The pay-off of each reaction is unequally spread between subpathways, similarly to that of glucose processing (respiration brings more energy units than fermentation) – 1:10, that is upstream reactions contribute one fitness unit for each molecule produced while downstream ones bring 10 units per reaction. Kinetic efficiencies are approximately half an order of magnitude higher than median values observed in datasets [Bar-Even et al., 2011], that is $k_f = 10^{6.25} M^{-1} s^{-1}$ and $k_{cat} = 10^{2.25} s^{-1}$ – considered as the default parameters throughout all this section (see Figure S12 for the influence of enzyme efficiencies). This echoes findings detailed in the main document about the relevance for cells to cope with permeability by allocating more to the part of the pathway downstream the metabolite that is subject to it, and, eventually, the promotion of CF interactions when degradation exceeds intermediate levels. When the pay-off is equally spread between subpathways, no CF emerges, either because shifting allocation is enough to cope with metabolite loss (with low degradation rates) or because the second subpathway cannot bring enough fitness (with high degradation rates) due to the loss – see Figure S8-B.

##### Text S3.1.2 Trait evolution plots: emergence of stable coalitions with high degradation rates

We then report the trait evolution plots – TEPs hereafter – that were used to build Figure 3 of the main paper. They illustrate that small mutational steps lead to the invasion of the singular strategy by a dimorphic population [Geritz et al., 1998, Brännström et al., 2013] and the further emergence of a protected cross-feeding coalition, where the second strain is specialised at processing the metabolite passively released by the first one – see Figure S9 for details. Notice that owing to computational difficulty, it is not possible to decide whether the singular strategy is stable for cases where fitnesses are closer from one another. Typically, in this situation, one would either expect that coalitions are favoured where the subtypes are in between the singular strategies and the specialist ones [Geritz et al., 1998], or that the outcome depends on the distribution of mutational effects.

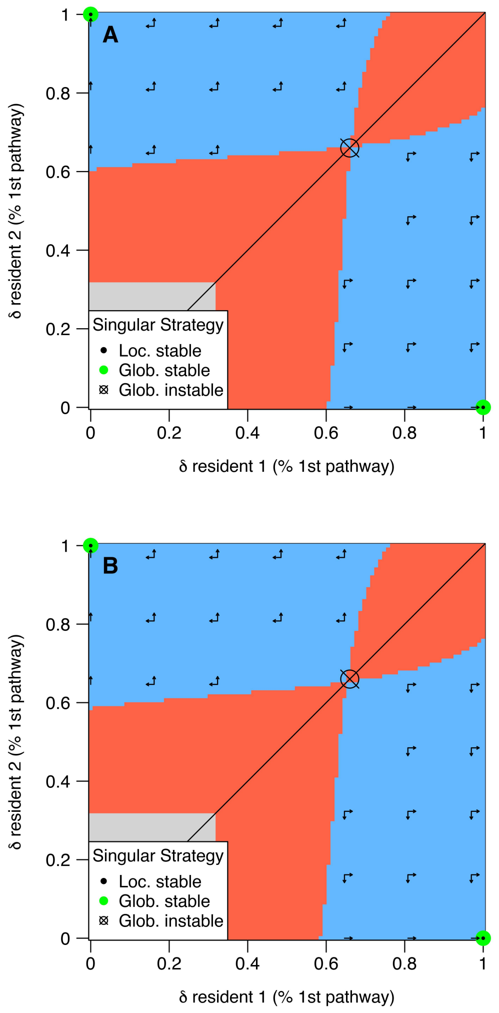


Figure S9: These TEPs show how a trait should evolve in a population comprised of two resident strategies when permeability equals $P = 10^{-5} dm \cdot s^{-1}$ in (A) and $P = 10^{-4} dm \cdot s^{-1}$ in (B). Each coalition is made up by two resident strategies, except on the bottom left toward upper right bisector for which the two resident strategies are identical. The red area is an area where coexistence is not possible, contrary to the blue one. In the blue area, we determine the fitness of each neighbouring mutant: there are four such mutants, except on the boundaries of the plot, as each resident can mutate and either increase or decrease its trait by one small unit. Here, we see that mutants that invade coalitions push the trait towards the upper left corner or the lower right one, which is exactly similar as these plots are symmetrical. As a consequence, the adaptive process should favour the emergence of CF interactions between one strain specialised at the transported nutrient and one strain specialised at processing the intermediate nutrient released by the previous one due to permeability.

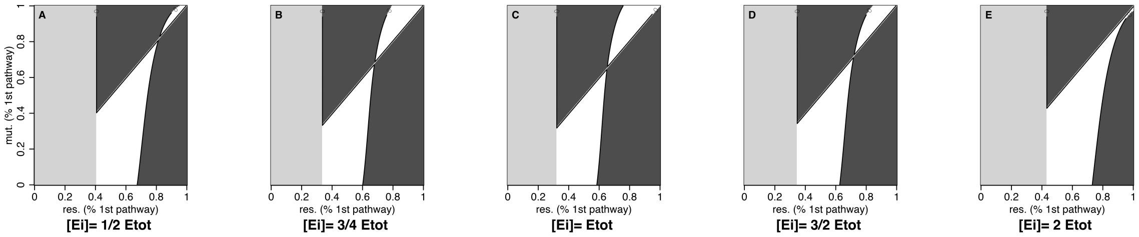


Figure S10: PIPs showing how the adapative process lead to mutual invasibility when $P = 10^{-4} dm \cdot s^{-1}$ no matter the level of the proteome fraction available for the focal pathway. (C) coincides with the case studied previously, where the fraction coincides with the adaptive value without permeability. On the left handside the total fraction is lower – (A): 1/2 and (B): 3/4 of the adaptive value without permeability – while on the right handside, it is higher – (D): 3/2 and (E): 2 of the same value. Besides, the area of mutual invasibility tends (predictably) to increase in both directions.

##### Text S3.1.3 Influence of the total available proteome fraction

In this section, we check that outcomes are not just a mere product of setting the total concentration to a given value. As to do so, we examined how PIPs are influenced by changes in the proteome fraction to allocate – see Figure S10 for $P = 10^{-4} dm \cdot s^{-1}$ and S16 in Appendix for $P = 10^{-5} dm \cdot s^{-1}$. Any of these changes produce PIPs where the area of mutual invasibility is wider, which should in turn favour coalitions. We also checked that these PIPs indeed yield the emergence of cross-feeding using TEPs – results not reported here. This implies that the coevolution between the total available proteome fraction and the allocation between subpathways also leads to cross-feeding interactions, because at any stage in the adaptive process of total proteome fraction, coalitions are favoured over the singular strategy.

#### Text S3.2 Influence of transport co-regulation and enzyme efficiencies

Thus far, we have considered that transporters were expressed no matter the strategy used. In this section, while keeping all features similar to the model case, we relax this assumption to explore its influence on mutual invasibility. Accordingly, transporters are considered to be perfectly co-regulated with the pathway, such that their concentration is a function of $\delta$: there are fully expressed when $\delta = 1$ and switched off when $\delta = 0$. We also considered a cost to the expression of transporters so that transporters that uptake the initial resource account for a few percent of the total budget in line with estimates found in the literature [Darbani et al., 2018]. Notice that the cost of transporters was of few influence and results were similar without considering it.

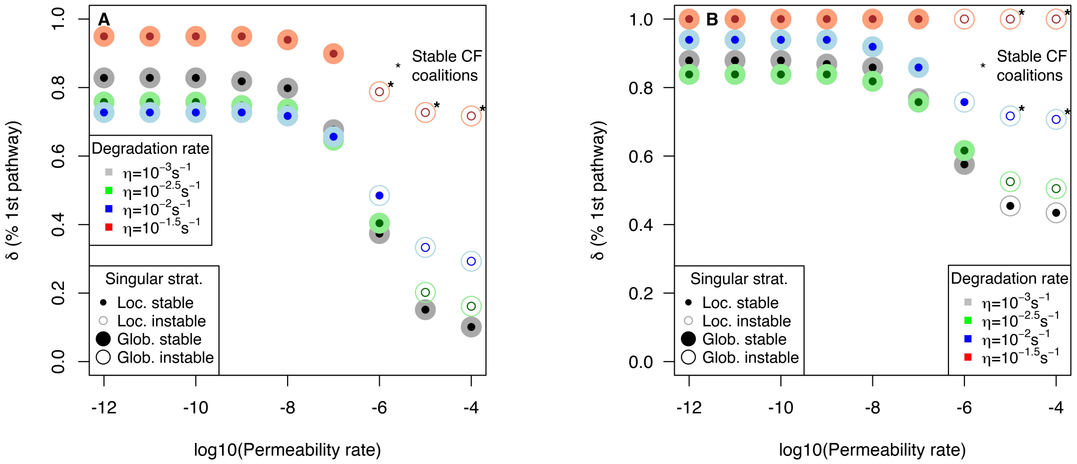


Figure S11: Influence of co-regulation between transporters and subpathways on the adaptive outcomes under the selective pressures of degradation and permeability: in (A), the pay-off is distributed according to the ratio 1:10 in favour of downstream reactions while in (B), it is similar in each subpathway. Including co-regulation favoured mutual invasibility, especially in the case of an equal pay-off along the whole pathway. Besides, it unlocks the possibility of CF coalitions – denoted with the star when TEPs always lead to this outcome – to a wider set of conditions, both in terms of degradation and permeability rates.

##### Text S3.2.1 Co-regulation of pathways and transporters facilitate the emergence of cross-feeding

Predictably, accounting for co-regulation (and cost) of transporters facilitates the emergence of mutual invasibility and cross-feeding interactions insofar as it both increases the relevance of focusing on the first sub-pathway and decreases that of keeping a part of the first sub-pathway when being partly specialised at processing the intermediate metabolite – see Figure S11 for more details.

##### Text S3.2.2 Enzyme efficiency also matters in the emergence of CF

Using the same framework with transporters co-regulation (only with the pay-off set to 1:10 in favour of downstream reactions), we then tested how enzyme efficiency would change the picture of adaptive outcomes. Like co-regulation, this parameter can facilitate or on the contrary complicate the advent of mutual invasibility and protected dimorphism (including CF) – see Figure S12.

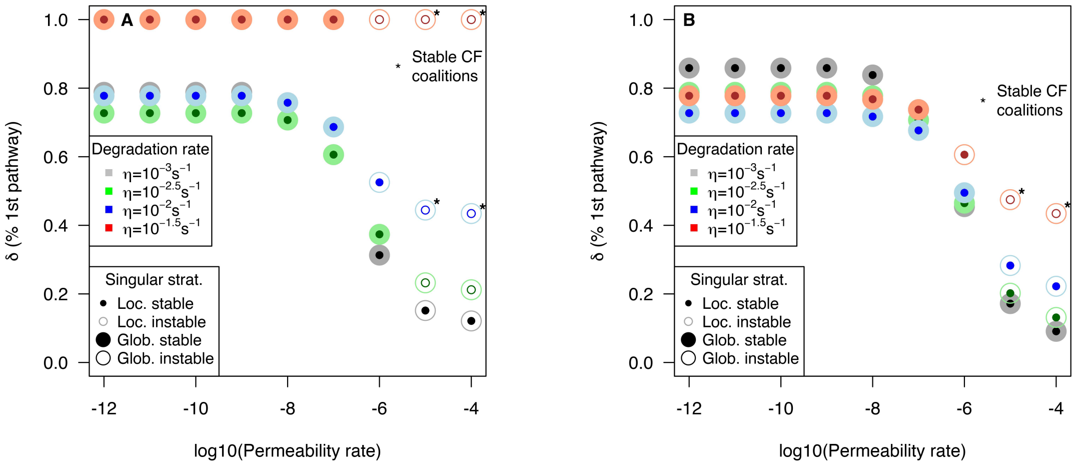


Figure S12: Influence of enzyme efficiencies on the adaptive outcomes under the selective pressures of degradation and permeability: in (A), the efficiency is close to the median found in datasets with kinetic parameters set to $k_f = 10^6 M^{-1}s^{-1}$ and $k_{cat} = 10^2 s^{-1}$ while in (B), they are nearly one order of magnitude higher with $k_f = 10^{6.5} M^{-1}s^{-1}$ and $k_{cat} = 10^{2.5} s^{-1}$. High kinetic efficiencies (B) narrows down the conditions in which mutual invasibility and CF interactions prevail whereas low ones (A) tend to promote their emergence – notice that for this enzyme efficiency, it was not possible to use the adaptive concentration without permeability for the highest degradation rate so that it had to be set manually, which slightly biased results (and did not enable us to determine what happens on the TEP with $P = 10^{-6} dm^{-1} \cdot s^{-1}$).

This parameter should play a part in the combination of conditions that leads to CF. Yet, it is harder to optimise efficiently as a response to a global constraint since its optimisation, besides being more difficult than for the level of expression, needs be synchronised with possibly all the other enzymes that are contributing to the pathway. For the sake of simplicity, we here considered that all enzymes have the same kinetic efficiency, while this latter is in fact the product of a tight combination of mechanistic constraints that may differ from one reaction to the next [Labourel and Rajon, 2021]. This seems nonetheless a reasonable assumption to make the case for permeability as a driver of CF since how the proteome is spread within each subpathway does not change the global impact they have on the constrained available proteome fraction: in other words, it is of few importance to know if the fraction dedicated to a pathway is equally spread or, on the contrary, heterogeneous among reactions, as long as the global impact on the subpathway is similar. What may still have an influence is the difference between the needs and costs of each subpathway, as seems to be the case for glycolysis and respiration [Basan et al., 2015].

#### Text S3.3 Toxicity and reversibility may also yield cross-feeding interactions

Hitherto, we have considered that all the internal constraints acting on reactions could be summarised through the degradation rate. As a proof of principle that it should not change much the predictions in terms of the critical CF-triggering permeability, we relax this assumption to show that accounting for the full set of mechanistic constraints can also favour the emergence of CF interactions.

We first tested how metabolite toxicity influences the outcomes by setting $\eta = 10^{-3}s^{-1}$ as to limit the impact of this parameter and studied three toxicity levels: $T = 10^{-2}M$, $T = 10^{-1}M$ and $T = 1M$. Enzyme kinetic efficiencies are set to $k_f = 10^{6.25}M^{-1}s^{-1}$ and $k_{cat} = 10^{2.25}s^{-1}$ and the pay-off is set to 1 : 10. Other parameters remain unchanged. Although slightly less restricted in terms of conditions, mutual invasibility arises with high levels of permeability but require toxicity to be at least moderate $T = 10^{-1}M$ – see Figure S13.

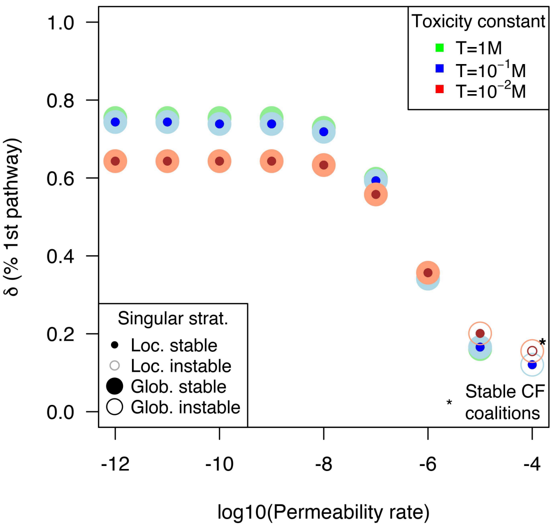


Figure S13: Influence of metabolite toxicity levels on the adaptive outcomes under the selective pressures of low degradation – $\eta = 10^{-3}s^{-1}$ – and permeability. Kinetic efficiencies are those of the model case – $k_f = 10^{6.25}M^{-1}s^{-1}$ and $k_{cat} = 10^{2.25}s^{-1}$. Mutual invasibility and stable CF still arises although they are respectively restricted to $P > 10^{-5}dm \cdot s^{-1}$ and $P > 10^{-4}dm \cdot s^{-1}$ and also require toxicity not to be too low. The main difference – not visible on this plot – is that CF involves a generalist strategy and a specialist cross-feeder rather than two specialist strategies. Notice that this case is very conservative because we did not consider the coregulation and cost of transporters, which proved to favour CF – see previous section.

The outcomes are sligthly different although branching points still emerge (see Figure S13), and, in some cases, cross-feeders may eventually have the edge on generalist strategies, leading to stable CF coalitions. Because there is little loss of metabolites, the generalist strategy has no interest to sacrifice its second subpathway and decreases as much as possible its investment in the first one. This looks like a Black Queen coexistence where it may seem costly to keep the first subpathway but it is nonetheless essential for the community to survive.

Finally, we report thereafter the influence of the membrane leakage when assuming the existence of reversibility within the pathway. In these cases again, cross-feeding interactions only arise when permeability reaches very high values $P > 10^{-6}dm \cdot s^{-1}$ – see Figure S14–(C-D), where toxicity is relatively high ($T = 10^{-1.5}M$). Although how reversibility is spread within reactions changes the emergence of CF with the two levels of toxicity considered on Figure S14, we checked that it is mainly affecting the tipping toxicity point (and more generally, the critical point for the combination of parameters shown previously to have an influence) where this change of adaptive behaviour occurs, as shown on Figure S17 of APPENDIX – where $T = 10^{-2}M$.

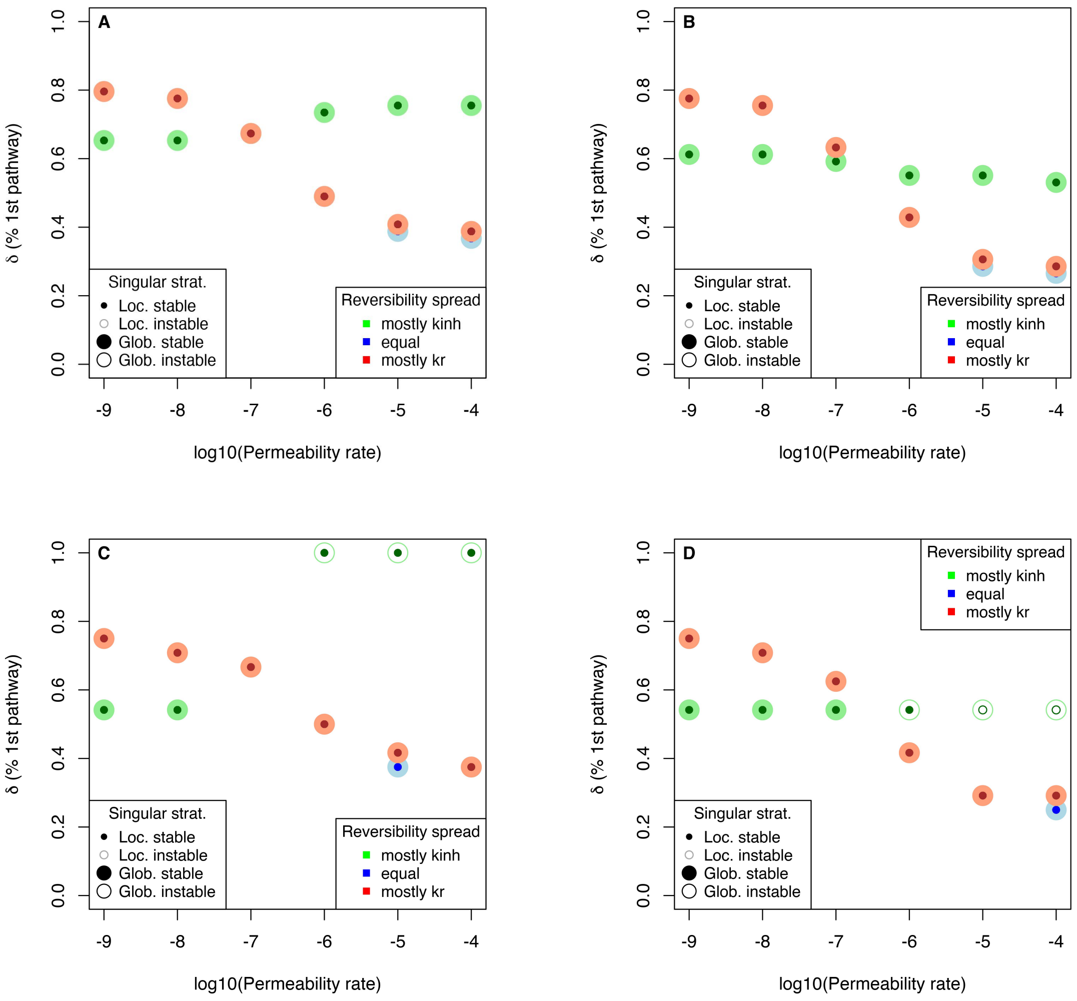


Figure S14: Influence of toxicity and reversibility on the adaptive outcomes when considering a low degradation rate $\eta = 10^{-3}s^{-1}$ and moderately high enzyme efficiencies corresponding to the model case $k_f = 10^{6.25}M^{-1}s^{-1}$ and $k_{cat} = 10^{2.25}s^{-1}$. Columns coincide to a constant pay-off value: the left handside – (A) and (C) – coincides with 1 : 1, while the right handside – (B) and (D) – coincides with 1 : 10. Rows coincide with two levels of toxicity: the upper row displays results for a moderately low toxicity $T = 10^{-0.5}M$ while the lower one shows those for a moderately high one $T = 10^{-1.5}M$. Reversibility has little impact on the adaptive outcomes as low toxicity still leads to the absence of mutual invasibility no matter the level of permeability. Yet, a higher toxicity triggers mutual invasibility only when reversibility is concentrated on $k_{inh}$ – see equation (S5) for details about how reversibility works. Reversibility thus slightly changes the conditions in which CF may be favoured (although with $T = 10^{-2}M$ – see Appendix – the discrepancy partly vanishes). One surprising phenomenon is that when reversibility influences mostly $k_{inh}$, this favours an overexpression of the first subpathway – see text for explanations about that. Notice that it was not possible to check what happens where mutual invasibility prevails owing to numerical inconsistencies due to reversibility.

Therefore, the conclusion that (only) high permeability rates – higher than $P = 10^{-7}dm\dot{}s^{-1}$ – can foster the evolution of cross-feeding interactions is robust across a wide range of conditions.

#### APPENDIX - Subset of plots complementing findings of the SM

Below, we first report results obtained when using the kinetic efficiencies of the main document and the last SM section:

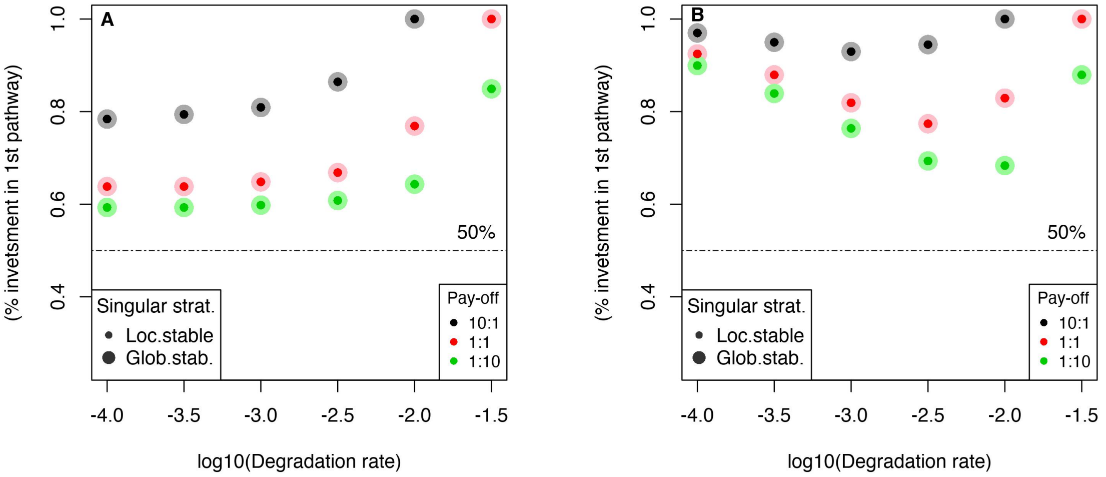


Figure S15: Counterpart of Figure S5 when kinetic parameters are those used in the model case with permeability, that is $k_f = 10^{6.25}M^{-1}s^{-1}$ and $k_{cat} = 10^{2.25}s^{-1}$. Selective constraints along an irreversible metabolic pathway push cells to focus on upstream reactions, no matter how energy gains are spread along the pathway – 10:1 as pay-off means that each reaction of the first subpathway provides 10 times the energy provided by each of the downstream reactions, and the other way around for 1:10. In (A), the influence of transporters is set aside by assuming a non-evolvable first enzyme concentration. Self-evidently, the degradation rate enhances the relevance to prioritise the first subpathway, because metabolites are progressively lost so that fewer metabolites can be processed downstream. More interestingly, even when the degradation rate is so low that very few metabolites are lost – see low degradation rates such as $\eta = 10^{-4}s^{-1}$ – it remains relevant to focus on the first subpathway (see text and next subsection for more details on this phenomenon). In (B), besides the two phenomenon described in (A), the impact on the transporter is shown to complexify the picture without changing the qualitative conclusion of an adaptive overexpression in upstream reactions: with low degradation rates, the transporter becomes the main constraint and because the selective pressure is low elsewhere in the pathway, it favors a very high overexpression of upstream reactions that tends to get lower with intermediate degradation rates because the seective pressure is homogenised along the pathway. Eventually, the loss of metabolites becomes so high that cells again should waive any downstream investment.

We then report the PIPs showing the effect of changing the total proteome fraction available to the focal pathway when $P = 10^{-5}dm \cdot s^{-1}$.

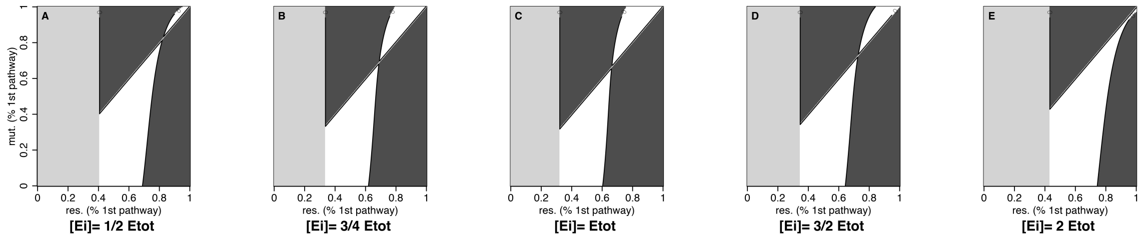


Figure S16: PIPs showing how the adapative process lead to mutual invasibility when $P = 10^{-5}dm \cdot s^{-1}$ no matter the level of the proteome fraction available for the focal pathway. (C) coincides with the model case, where the fraction coincides with the adaptive value without permeability. On the left handside the total fraction is lower – (A): 1/2 and (B): 3/4 of the adaptive value without permeability – while on the right handside, it is higher – (D): 3/2 and (E): 2 of the same value. Besides, the area of mutual invasibility tends (predictably) to increase in both directions.

Finally, we report the outcomes of proteome allocation with a high toxicity rate $T = 10^{-2}M$ and reversibility as to show that branching points may exist no matter how reversibility is spread within a reaction.

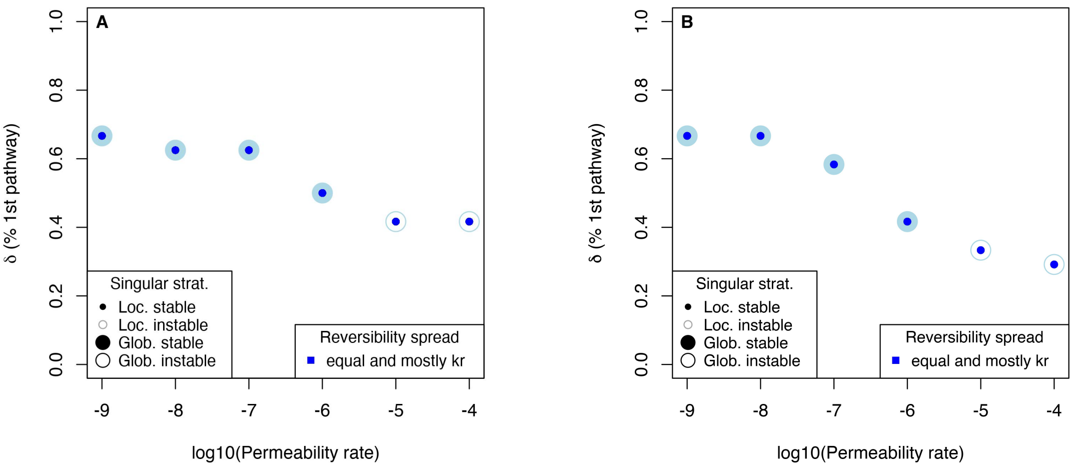


Figure S17: Influence of toxicity and reversibility on the adaptive outcomes when considering a low degradation rate $\eta = 10^{-3}s^{-1}$, a high toxicity rate $T = 10^{-2}M$ and moderately high enzyme efficiencies corresponding to the model case $k_f = 10^{6.25}M^{-1}s^{-1}$ and $k_{cat} = 10^{2.25}s^{-1}$. As described in the last section of the SM, global instability do emerge no matter how the reversibility is spread provided the level of toxicity is high enough and permeability exceeds the same highlighted critical point than for other conditions (around $P = 10^{-6}dm \cdot s^{-1}$). Two different pay-off distributions are considered (A):1 : 1 and (B):1 : 10, without any remarkable influence on the results; the way reversibility is spread is also of little influence.
